# Evaluating phasic transcutaneous vagus nerve stimulation (taVNS) with pupil dilation: the importance of stimulation intensity and sensory perception

**DOI:** 10.1101/2024.07.27.605407

**Authors:** Mareike Ludwig, Calida Pereira, Marius Keute, Emrah Düzel, Matthew J. Betts, Dorothea Hämmerer

## Abstract

The efficacy of transcutaneous auricular vagus nerve stimulation (taVNS) as a non-invasive method to modulate physiological markers of noradrenergic activity of the Locus Coeruleus (LC), such as pupil dilation, is increasingly more discussed. However, taVNS studies show high heterogeneity of stimulation effects. Therefore, a taVNS setup was established here to test different frequencies (**10 Hz** and **25 Hz**) and intensities (**3 mA** and **5 mA**) during phasic stimulation (**3 s**) with time-synchronous recording of pupil dilation in younger adults. Specifically, phasic real taVNS and higher intensity led to increased pupil dilation, which is consistent with phasic invasive VNS studies in animals. The results also suggest that the influence of intensity on pupil dilation may be stronger than that of frequency. However, there was an attenuation of taVNS-induced pupil dilation when differences in perception of sensations were considered. Specifically, pupil dilation during phasic stimulation increased with perceived stimulation intensity. The extent to which the effect of taVNS induces pupil dilation and the involvement of sensory perception in the stimulation process are discussed here and require more extensive research. Additionally, it is crucial to strive for comparable stimulation sensations during systematic parameter testing in order to investigate possible effects of phasic taVNS on pupil dilation in more detail.

## Introduction

The efficacy of transcutaneous auricular vagus nerve stimulation (taVNS) as a non-invasive method for modulating the noradrenergic system of the locus coeruleus (LC-NE system) is increasingly being discussed^1–3^. TaVNS modulation occurs by the transmission of excitation from the Auricular Branch of the Vagus Nerve (ABVN) through the remaining nerve fibre bundles of the vagus nerve via the nucleus tractus solitarius (NTS) to the LC, both of which are located in the brainstem ^4,5^. The LC is the main source of NE in the brain and its activation can be either in a “*tonic”* (continuous activity: maintaining arousal and attention) or “*phasic”* mode (rapid bursts: brief changes in attention and arousal due to for example salient or novel stimuli), which is linked to distinct levels of NE release^6–11^. Since animal studies have shown that *phasic LC stimulation* causes an increase in *pupil dilation*^12,13^, and because both animal and human studies have shown that pupil dilation covaries with both spontaneous LC activity^11,14^ and transient increases in LC activity (such as with task-relevant or salient factors)^15–17^, changes in pupil dilation can serve as an indicator of LC-NE activity. However, this link is not exclusive, as other brain structures (e.g., hypothalamus, superior colliculus) can also cause pupil dilation^12,18^, and noradrenergic and cholinergic axons are both involved^13,19,20^. It has also been shown that iVNS can modulate neurotransmitters such as acetylcholine and dopamine in rats^21–23^. Changes in pupil dilation can thus be considered as an *indirect outcome measure* to investigate the effects of taVNS.

Using invasive VNS (iVNS), it has been shown that increased LC firing rate could be achieved by adjusting stimulation parameters, such as higher intensities and longer pulse widths during phasic stimulation^22–24^. In particular, experimental testing of various stimulation parameters revealed more dilated pupils at higher stimulation parameters in animals (for review see Ludwig et al.^1^). While iVNS is employed in human subjects as well, taVNS is more extensively utilized and has a less intricate therapeutic application. However, the effectiveness of taVNS studies to date has been characterized by high heterogeneity and low reliability of stimulation effects (see Farmer at al.^25^ & Ludwig et al.^1^ for review). A promising study in humans stimulated with brief bursts (3.4 s) demonstrated increased pupil dilation^3^, which is consistent with the results reported above from animal research^24^. These results, keeping the stimulation parameters and protocol the same as in Sharon et al.^3^, could also be replicated by Lloyed et al.^26^ with comparable intensities during taVNS (2.3 ± 1.3 mA). In most human taVNS studies (see Burger et al.^27^, Farmer at al.^25^ & Ludwig et al.^1^ for review), an individual stimulation intensity below the respective pain threshold was applied to control for the sensory effects between subjects and between real vs. sham stimulation within the same subject. A recent taVNS study systematically tested different pulse widths (200, 400 μs) with different intensities during phasic stimulation (5 s), while with increased stimulation intensity and pulse width, pupil dilated more during taVNS as compared to sham stimulation^28^. However, it remains unclear which parameter combination under which comparable stimulation sensations per stimulation location leads to the increased pupil dilation. When interpreting the significance of pupil dilation in i/taVNS, it is therefore important to consider the different anatomical structures and pathways as well as other possible influencing factors, such as the perception of the stimulation sensations and the stimulation parameters.

In the present study, a taVNS setup was established to allow systematic testing of different stimulation parameters with time-synchronous recording of pupil dilation by comparing phasic real and sham stimulation within healthy younger adults on a single day. Specifically, in a randomized and counterbalanced order within subjects, **10 Hz** and **25 Hz** frequency were tested in combination with **3 mA** and **5 mA** intensity while subjects looked at a fixation cross during **3 s of phasic stimulation**. The comparison between the frequency 25 Hz and 10 Hz was chosen in agreement with most studies applying 25 Hz (see Table 1 & 2 Farmer et al.^25^) which has also been shown to induce LC activation in taVNS studies in humans^2,29^, where the 10 Hz served as a lower frequency. The comparison between an intensity of 3 mA and 5 mA was chosen to apply findings from invasive animal studies to the non-invasive approach used here, as animal studies have shown that stronger intensities result in greater phasic LC activity^22–24^, whereas a threshold of 2.5 mA showed the highest firing rate of LC neurons^24^. The 5 mA intensity was the maximum that could be set with the stimulator, and 3 mA served accordingly as a lower intensity for comparison. Additionally, the study investigated the extent to which the effect of taVNS alone led to pupil dilation and the extent to which perception of sensation was involved in the stimulation process, as this has not yet been investigated in detail in previous i/taVNS studies.

In line with greater pupil dilation during phasic real i/taVNS^3,22–24,28^, we expected (**1**) greater pupil dilation during phasic real compared to sham stimulation. Furthermore, since iVNS studies showed greater maximal LC discharge rate and greater pupil dilation during higher intensity and frequency^22–24^, we expected greater pupil dilation during (**2**) higher intensity and (**3**) higher frequency stimulation. Similarly, we expected that the pupil dilation would (**4**) gradually contract over time^3,22,23^. Given the potential influence of subjective sensations of stimulation on pupil dilation, this was considered and discussed accordingly.

## Results

### Model comparisons

A distinct Linear Mixed Model (LMM) was fitted for each time window of analysis, that is **(I)** the 3 s during ‘**on stimulation’** time window, **(II)** the first 3 s during off stimulation ‘**immediate response**’ time window and **(III)** the subsequent last 10 s during off stimulation ‘**delayed response**’ time window using the same criteria (see Methods; see Supplementary Table S1-S4).

First, to evaluate the effects of different stimulation conditions, intensities and frequencies on pupil dilation, the forward model selection considered variables for ***‘stimulation’*** *[real (1) vs. sham (0)]*, ***‘frequency’*** *[high (1) vs. low (0)]* and ***‘intensity’*** *[high (1) vs. low (0)])*. When testing whether interactions between stimulation, frequency and intensity further improved the model, the results for each distinct model at each time window (I-III) showed no significant improvement over the best model without interactions (model m_4: StimIntFreq-LMM) with the following lowest AIC values in the model comparison for (I) AIC = 29404 (χ^2^ = 102.95, p < 0.001), (II) AIC = 37688 (χ^2^ = 76.54, p < 0.001) and (III) AIC = 39305 (χ^2^ = 8.27, p = 0.004):

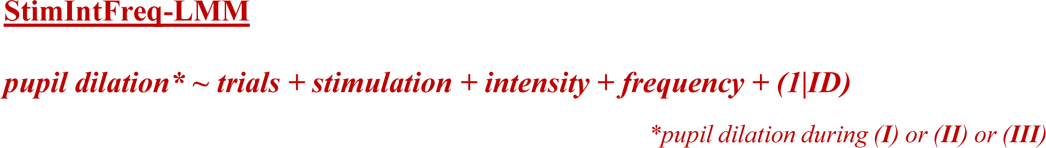

Second, based on the StimIntFreq-LMM model, further factors which can modulate or mediate stimulation effects on pupil dilations were added (see Methods; see Supplementary Fig. S1 & Table S1). The following factors were added stepwise: ***‘VAS’*** (subjective perception of sensations), ***‘sensitivity’*** *[sensitive (1) vs. not sensitive (0)], **‘real_first’** [counterbalanced: real (1) before sham (0) stimulation], **‘position’** (four different stimulation combination possibilities),**’gender’** [female (1) vs. male (0),**’sporty’** [sporty (1) vs. non-sport (0)]:* The renewed model comparison now showed that the model with VAS (model m_4_1: StimIntFreq-VAS-LMM) was the best model at each time window (I-III) with the following lowest AIC values in the model comparison for (I) AIC = 29367 (χ^2^ = 38.76, p < 0.001), (II) AIC = 37659 (χ^2^ = 30.81, p < 0.001).

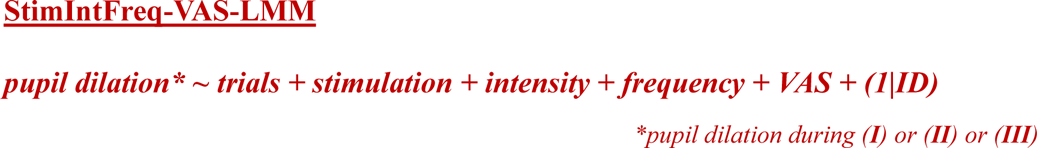

However, for (III) **‘delayed response’** the StimIntFreq-LMM was not significant (AIC = 39304 (χ^2^ = 3.32, p = 0.07)), but StimIntFreq-LMM was again the best model (AIC = 39305 (χ^2^ = 8.27, p = 0.004)) (see Supplementary Table S4).

An exploratory analysis (see Supplementary Figure S4) was added to investigate potential interactions between VAS and stimulation condition and parameters in influencing pupil size controlled for sensitivity.

### Increased pupil dilation during phasic real taVNS? Exploring the impact of subjective perception of sensations

In accordance with increased pupil dilation during phasic i/taVNS stimulation^3,22,28^, the **StimIntFreq-LMM** model suggested that pupil dilation was increased during real (M±SD: 0.18±0.03) as compared to sham (M±SD: 0.1±0.03) stimulation during **(I)** ‘on stimulation’ (χ^2^ = 18.98, p < 0.001). Additionally, during the (**II**) ‘immediate response’, immediately after stimulation was turned off, pupil dilation was still increased during real (M±SD: 0.15±0.04) as compared to sham (M±SD: 0.04±0.04) stimulation (χ^2^ = 15.99, p < 0.001), while there was no significant difference between real (M±SD: −0.04±0.02) and sham (M±SD: −0.06±0.02) stimulation (χ^2^ = 0.61, p = 0.43) for the (**III**) ‘delayed response’, 3 s after stimulation was turned off (see Fig. 1; Supplementary Table S5). However, the **StimIntFreq-VAS-LMM** model revealed that VAS explained significant proportion of pupil variance during **(I)** ‘on stimulation’ (χ^2^ = 38.79, p < 0.001) and **(II)** ‘immediate response’ (χ^2^ = 30.82, p < 0.001) ((III) ‘delayed response’ (χ^2^ = 3.37, p = 0.07)). Therefore, the difference between real and sham stimulation was no longer statistically explainable (Supplementary Table S6) during **(I)** ‘on stimulation’ (χ^2^ = 2.69, p = 0.1) and the **(II)** ‘immediate response’ (χ^2^ = 2.42, p = 0.12), after accounting for condition differences in VAS. This suggests that effects of real vs. sham stimulation and effects of different subjective perception of sensations of real vs. sham stimulation on pupil dilations cannot be distinguished statistically with the stimulation parameters employed here.

**Figure 1.**
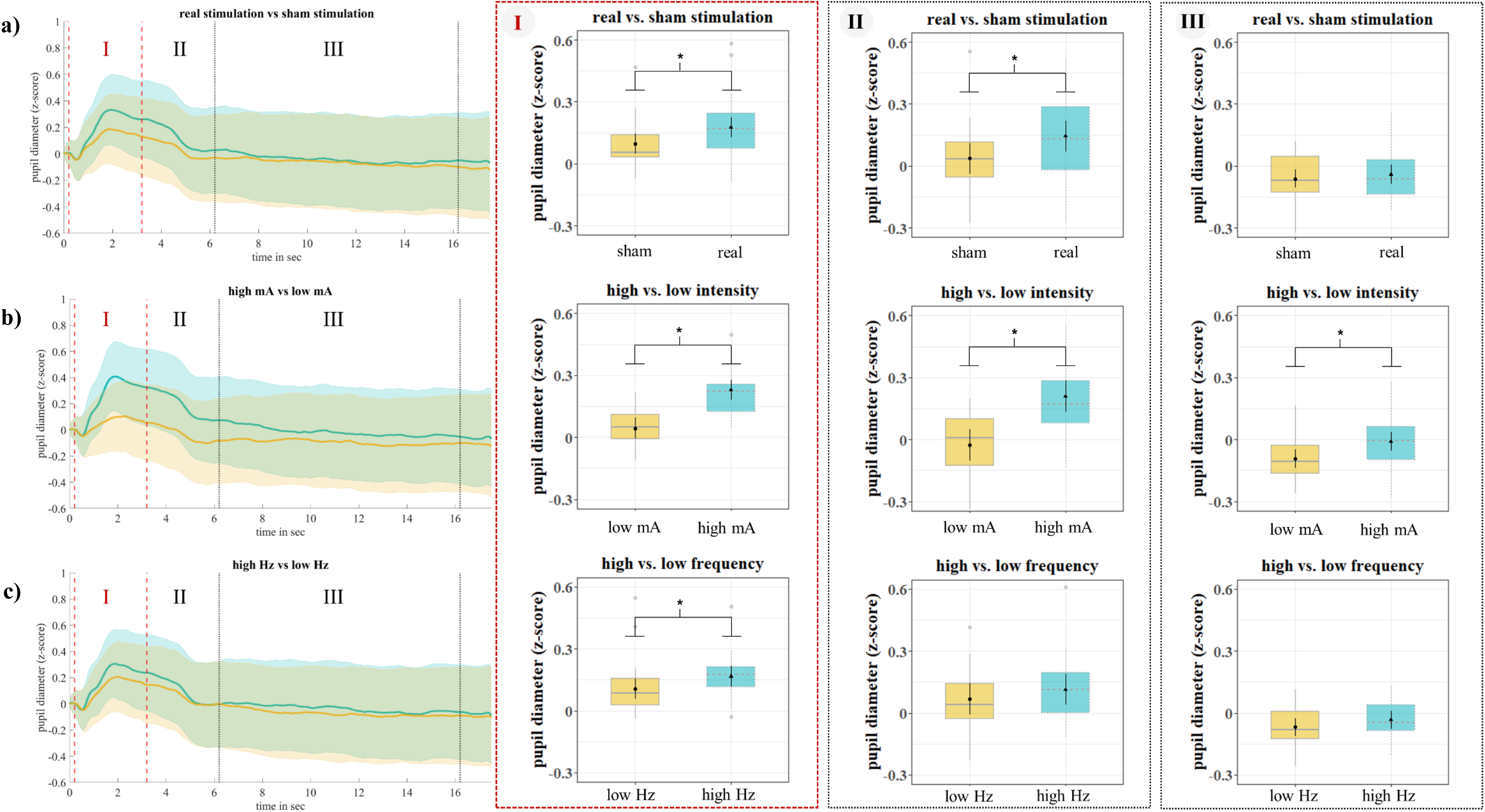
Changes in pupil dilation during real and sham stimulation for low and high stimulation intensity and frequency. Pupil diameters for **a)** real (turquoise) and sham (ochre) stimulation, **b)** high (turquoise) and low (ochre) intensity and **c)** high (turquoise) and low (ochre) frequency. Shadowed lines represent the standard error across subjects (left panel). The dashed vertical red lines indicate the time window of (I) ‘**on stimulation’**, the (II) ‘**immediate response’** following to the first dashed vertical black line and the subsequent (III) ‘**delayed response’** to the second dashed vertical black line. The boxplots for the individual time windows are based on the ***StimIntFreq-LMM***, the asterisks indicate significant differences between conditions. The reader is encouraged to look at Supplementary Fig. S5 to see the influence of VAS on stimulation and stimulation parameters in the **StimIntFreq-VAS-LMM** model.

### The importance of high intensity stimulation and the impact of subjective perception of sensations

In line with iVNS approaches in animals showing increased pupil dilation during higher intensity stimulation^23,24^, **StimIntFreq-LMM** suggested that pupil dilation was increased during higher (M±SD: 0.23±0.03) as compared to lower (M±SD: 0.04±0.03) intensity during (**I**) ‘on stimulation’ (χ^2^ = 103.40, p < 0.001). During the **(II)** ‘immediate response’, pupil dilation was still increased during higher (M±SD: 0.21±0.04) as compared to lower (M±SD: - 0.03±0.04) intensity (χ^2^ = 76.79, p < 0.001), and there was still a significant difference between higher (M±SD: −0.01±0.02) and lower (M±SD: −0.09±0.02) intensity (χ^2^ = 8.26, p = 0.004) for the (**III**) ‘delayed response’ (see Fig. 1; Supplementary Table S5). When additionally controlling for VAS differences across stimulation conditions in the **StimIntFreq-VAS-LMM** model, pupil dilation was still increased during higher (M±SD: (I) 0.20±0.03; (II) 0.17±0.04) as compared to lower (M±SD: (I) 0.08±0.03; (II) 0.02±0.04) intensity during (**I**) ‘on stimulation’ (χ^2^ = 28.53, p < 0.001), and during the **(II)** ‘immediate response’ (χ^2^ = 20.10, p < 0.001), but not during **(III) ‘**delayed response’ (χ^2^ = 2.76, p = 0.1) (see Supplementary Table S6). This suggests that stimulation intensity, especially higher intensity levels, and subjective perception of stimulation intensity may contribute to pupil dilation to varying degrees.

### Increased pupil dilation during higher frequency only during phasic stimulation and the impact of subjective perception of sensations

The **StimIntFreq-LMM** suggested that pupil dilation was increased during higher (M±SD: 0.17±0.03) as compared to lower (M±SD: 0.11±0.03) frequency during (**I**) ‘on stimulation’ (χ^2^ = 10.88, p = 0.001). During the **(II)** ‘immediate response’, pupil dilation was statistically not increased during higher (M±SD: 0.11±0.04) as compared to lower (M±SD: 0.07±0.04) frequency (χ^2^ = 2.92, p = 0.09), and there was also no significant difference between higher (M±SD: −0.03±0.02) and lower (M±SD: - 0.07±0.02) frequency (χ^2^ = 1.53, p = 0.22) anymore for the (**III**) ‘delayed response’ (see Fig. 1; Supplementary Table S5). These results would be in line with iVNS approaches in animals showing increased pupil dilation during higher frequency stimulation over a shorter period of time (Hulsey et al., 2019). However, after controlling for subjective perception of sensations in the **StimIntFreq-VAS-LMM** model, there was no effect of different stimulation frequencies observable neither for (**I**) ‘on stimulation’ (χ^2^ = 2.85, p = 0.1), nor for **(II)** ‘immediate response’ (χ^2^ = 0.09, p = 0.76) and **(III) ‘**delayed response’ (χ^2^ = 0.68, p = 0.41) (see Supplementary Table S6).

Since VAS was not kept constant due to systematically testing of different stimulation parameters and given that **StimIntFreq-VAS-LMM** (the most appropriate model given the data) suggested including VAS as an explanatory variable for pupil dilation, it was necessary to evaluate the extent to which the subjectively experienced sensory effects are statistically associated with the stimulation parameters and changes in pupil dilation.

### Subjective higher perception of sensation (VAS) in response to stimulation

The subjective perception of sensations (VAS) was not only higher for **(1)** *real stimulation* (M = 4.40, SD = 0.34) compared to sham stimulation (M = 2.97, SD = 0.29), (F(1,20) = 17.85, p < 0.001), but also for **(2)** *high frequency* (M = 4.08, SD = 0.28) compared to low frequency (M = 3.29, SD = 0.28), (F(1,20) = 19.57, p < 0.001), and **(3)** *high intensity* (M = 4.56, SD = 0.23) compared to low intensity (M = 2.81, SD = 0.23), (F(1,20) = 70.90, p < 0.001). There was no significant effect for either gender (F(1,20) = 0.02, p = 0.88), sporty (F(1,20) = 0.17, p = 0.69), or sensitivity (F(1,20) = 0.07, p = 0.79). There was a significant interaction between *sensitivity and stimulation* (F(1,20) = 4.86, p = 0.04), whereas sensitive subjects perceived higher sensations during real stimulation (M = 4.71, SD = 0.59) than during sham stimulation (M = 2.51, SD = 0.5); t(20) = 3.78, p = 0.006. Additionally there were trends for interactions between (1) stimulation and frequency (F(1,20) = 4.06, p = 0.06) as well as between (2) sensitivity and frequency (F(1,20) = 3.61, p = 0.07). However, there was no significant interaction between in stimulation and intensity (F(1,20) = 0.20, p = 0.66) (see Supplementary Fig. S3).

### Relationship between subjective perception of sensations (VAS) of stimulation and pupil dilation across subjects

To further investigate the potential effects of VAS on pupil dilation across subjects, VAS after each stimulation session and corresponding pupil dilation (averaged per subject across all trials within a stimulation condition) were correlated (see Supplementary Fig. S2, Supplementary Table S7). Correlations between VAS and pupil dilation considering outlier correction (see methods) were found for **a)** real stimulation: low intensity & low frequency (r = 0.43, p = 0.04), **f)** sham stimulation: high intensity and low frequency (r = 0.52, p = 0.01) as well as **g)** sham stimulation: low intensity & high frequency (r = - 0.45, p = 0.04). However, there were no correlation between VAS and pupil dilation for **b)** real stimulation: high intensity & low frequency (r = 0.01, p = 0.95), **c)** real stimulation: low intensity & high frequency (r = −0.11, p = 0.62), **d)** real stimulation: high intensity & high frequency (r = 0.31, p = 0.14), **e)** sham stimulation: low intensity & low frequency (r = 0.19, p = 0.37), **h)** sham stimulation: high intensity & high frequency (r = 0.27, p = 0.20). Thus, while higher mean pupil dilation was not consistently associated with higher VAS ratings across all subjects, three instances of significant associations between pupil dilation and VAS (two positive, one negative) were observed, suggesting that interindividual differences in subjective perceptions of stimulation also add variance to pupil ratings of stimulation effects.

## Discussion

This study examined the effects of phasic taVNS on pupil dilation by systematically testing different frequencies (10 Hz vs. 25 Hz) and intensities (3 mA vs. 5 mA) within younger healthy subjects, while keeping pulse width and total duration of stimulation constant during a luminance-controlled resting state task. Due to the systematic testing of varied frequencies and intensities, it was not feasible in the present study to maintain consistent VAS ratings across stimulation conditions, which necessitated acquiring VAS ratings after each stimulation session. Subjective perception of sensations due to stimulation was higher for real than for sham stimulation, which is also consistent with a recent phasic taVNS study^28^, and at higher intensities and frequencies compared with lower ones. The effects of taVNS on pupil dilation were investigated not only based on different stimulation conditions and parameters but also regarding potential confounds in predicting pupil dilation related to subjective perceptions of sensations due to stimulation.

In line with prior i/taVNS studies, **phasic taVNS** led to increased pupil dilation during real compared to sham stimulation, at higher compared to lower intensity, and during higher compared to lower frequency stimulation (for review see Ludwig et al.^1^). In the present study, we also examined the temporal dynamics of these effects and the duration of their persistence following the cessation of phasic stimulation. The effects of the stimulation and stimulation parameters on the pupils were generally more pronounced during the (I) 3 s of phasic ‘on stimulation’ than in the (II) 3 seconds after stimulation (‘immediate response) and in a subsequent (III) 10-second time-window (‘delayed response’). Differential effects in the time course for the here evaluated different stimulation parameters were observed. While the differences in pupil dilation between real and sham stimulation persisted for another 3 seconds after the end of stimulation, differential effects of stimulation frequencies were no longer detectable after stimulation was turned off, whereas effects of stimulation intensity persisted in a time window of 6.2 to 16.2 s after stimulation was turned off.

In general, the time course of the increase and the peak of pupil dilations during the 3 s of **phasic real stimulation** as well as the decrease afterwards corresponds to the results of the 3.4 s phasic stimulation of Sharon et al.^3^ and their replication^26^. Similarly, a recent study using 5 s phasic stimulation showed increased pupil dilation during taVNS, which decreased shortly after the peak^28^. Another study also showed that far shorter phasic taVNS (∼ 600 ms) equally led to an increased pupil dilation with canal stimulation leading to larger pupil dilation than conchae stimulation compared to sham stimulation^30^. The observed increase in pupil dilation during phasic taVNS may be consistent with previous findings in animal studies, which have demonstrated increased LC activation and increased NE release with phasic stimulation^10,24^. Importantly, the best-fitting model ‘StimIntFreq-VAS-LMM’ revealed that differences in the **subjective perception of sensations** due to stimulation evaluated here explained a significant proportion of the stimulation effects on pupil dilation. In particular, the difference between real and sham stimulation could no longer be explained statistically, but also differences in stimulation frequencies and intensities were almost fully accounted for. This may imply that differences in the subjective perception of real vs. sham stimulation are either exclusive drivers of pupil differences between real and sham stimulation and differences between low and high intensity and frequency or that effects of stimulation and perception of the stimulation overlap to a statistically indistinguishable amount with the given stimulation parameters used here. The latter explanation could be considered more likely as Sharon et al.^3^ showed pupil dilation due to phasic real stimulation compared to sham at constant VAS.

The present results may suggest that the level of **intensity**, particularly at higher levels, plays a crucial role in the stimulation process and its possible impact on pupil dilation. Indeed, higher intensities (0-2.5 mA) and longer pulse widths (0-500 μs) during iVNS in rats did modulate LC activation^24^, while increased pulse width (100, 200, 400, 800 μs) during iVNS led to an increased pupil dilation^22^. D’Agostini et al.^28^ also indicated a more dilated pupil by increasing the pulse width (200 or 400 μs) with intensity (.2,.5, calibration intensity: 1.19 ±.65 mA (taVNS) and 1.49 ±.73 mA (sham)), which may suggest that pulse width appears to be an important contributing factor. Since a wider pulse width (500 μs) is thought to result in a narrower range of VNS intensities^31^, it may explain why D’Agositini et al.^28^ were able to report pupil dilation during phasic tAVNS at very low intensities due to a higher pulse width. It may also explain why subjects in this study were able to tolerate up to 5 mA during the short 3 s phasic stimulation with the moderate 250 μs pulse width. Additionally, **frequency** seems to have a less strong influence on the pupil. Thus, iVNS studies showed increased pupil dilation during stimulation at 20 Hz compared to 10 Hz and 5 Hz^22^ and a greater increase in LC firing rate over a shorter period of time during high frequencies (constant parameters: 0.8 mA, 100 μs, 16 pulses and tested frequencies of 0, 7.5, 15, 30, 60, 120 Hz), but that higher frequency did not affect the total amount or neuronal activity in the LC^24^. Nevertheless, since the stimulation intensity and frequency effects occurred in the absence of a real vs. sham stimulation effect (based on limited sample size (N=24)), it is difficult to assume that it reflects effects mediated by the vagus nerve. An additional process affecting pupil dilation, such as attention-induced modulation of pupil size, might explain this effect^32,33^.

Therefore, when using pupillometry as an outcome measure for taVNS studies, different levels of subjective perception of sensations between stimulation conditions are a confounding factor. More importantly, as the typical locations for real and sham stimulation showed a higher VAS rating for real stimulation at the cymba conchae for the same stimulation parameters, it would be important to investigate to what extent alternative sham stimulation locations (e.g., earlobe, scapha) generally show a comparable level of stimulation induced sensations to alternative real stimulation locations (e.g., cymba conchae, tragus). A typical solution is to keep the sensations for real and sham stimulation constant^3,25,27^. The observation of potential greater effects of real stimulation under these conditions thus allows a conservative assessment of the benefits of real stimulation over sham stimulation. It is a major challenge to investigate the effects of different stimulation parameters under constant stimulation sensation, which is of utmost importance in human taVNS studies. An attempt to fulfil this requirement was shown, for example, in a taVNS study in humans in which not only pairs with lower frequencies and higher amplitudes were tested with pairs of higher frequencies with lower amplitudes, but also during respiratory auricular vagal afferent nerve stimulation (RAVANS), controlling for subjective perception of stimulation^29^. Especially RAVANS enables the stimulation at the same location during inhalation and exhalation. Both 100 Hz and 2 Hz led to increased LC activation, while no stimulation was applied during sham stimulation^29^. Considering that the LC is involved in attentional processes^34^ and studies in monkeys have already shown that there is increased phasic LC activation during discriminative tasks^35^, the question arises whether 2 and 100 Hz were more salient and could be discriminated, contributing to increased LC activation during RAVANS; potentially suggesting the influence of attention-induced modulation during taVNS. Additionally, even if the subjective perception of sensations due to stimulation is kept constant, sensory features that allow discrimination of different stimulation parameters could influence pupillometric responses to taVNS. It could be investigated whether subjects can **discriminate** different qualities of stimulation parameters (e.g., increasing intensity with constant pulse width or vice versa) and whether certain parameter combinations are more salient. It is assumed that the stimulation activates A, B and C fibres of the cervical vagus nerve (cVNS) to varying degrees^36,37^, while tAVNS may have been transmitted by thick afferent A-beta axons, as discussed by Safi et al.^38^. Likewise, an iVNS in rats in which vagal afferent C-fibres were destroyed with capsaicin still showed reduced seizures^39^. Therefore, another option could be to apply **anaesthetic cream** (e.g., lidocaine) to the ear area to be stimulated to suppress the sensory perception of the stimulation. However, it has been shown that when nerves were exposed to lidocaine, A and B fibers were blocked^40^ and, additionally, firing rate of the vagus nerve also decreased when lidocaine was administered distal to the cervical vagus nerve^41^. Accordingly, with such anaesthetic creams, it cannot be ruled out that nerve fibre connections could be blocked, which could be important for the transmission of electrical impulses of tAVNS. Further experiments including invasive fibre recordings are needed to possibly determine optimal doses (probably the minimum most effective) of anaesthetic creams during taVNS in humans. Interestingly, it may also be possible to modulate the effects of stimulation parameters and the sensations of stimulations separately, as increasing firing rates and frequencies have been shown to have different effects on LC activity, with increasing frequencies decaying much earlier than increasing intensities^24^. As already discussed in a review^1^, it remains a central question which comparable real and sham stimulation locations and comparable stimulation sensations are best suited for different stimulation parameters. Limitation should be mentioned, as further research is needed to systematically investigate possible carry-over effects of single taVNS sessions with different durations on pupil dilation. It is currently unclear whether the duration of stimulation is related to the effects of taVNS on pupil dilation and which wash-out period should be chosen between sessions. As the subjects had already received real and sham stimulation in a counterbalanced order during the emotional memory task (approx. 44 min stimulation, see Fig. 2) before the resting state task, followed by wash-out of approx. 45-60 min, it is challenging to assess to what extent the stimulation effects observed here are partly due to a prior ‘pre-stimulation effect’. Future double-blinded studies are needed to determine whether this was the case and if so, to what extent. Moreover, our subject sample was relatively small (N=24), and while variance related to random slopes in intensity effects could be captured, we were not able to specify statistical models incorporating the full random effects structure due to model convergence issues. In order to obtain more robust model estimates and maximize generalizability, a larger sample size would be desirable for future studies.

**Figure 2.**
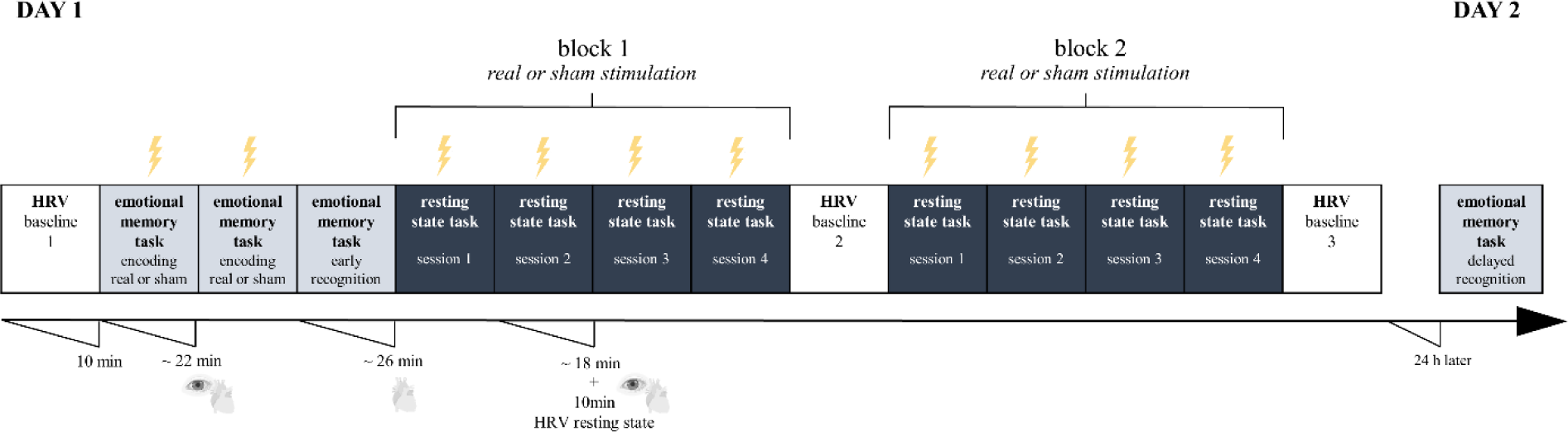
Study procedure overview. The study was conducted as a sham-controlled, single-blind, within-subject, counterbalanced, randomized design with a one-day stimulation protocol. During the emotional memory task real or sham stimulation was applied with highest stimulation parameters (5 mA, 25 Hz), while during the resting state task 4 different parameter combinations were systematically tested (3 mA and 5 mA with 10 Hz and 25 Hz) in block 1 compared to block 2 (real or sham stimulation). Additionally, heart-rate variability (HRV) as well as changes in pupil dilation during taVNS were recorded.

The present study demonstrates how crucial it is to consider the perception of sensations brought on by phasic taVNS, as a large proportion of studies either control for VAS or do not report systematically at all. Since VAS ratings are subjective and pre- and post-stimulation ratings are likely to capture potential habituation effects, more objective measures such as skin conductance should be included and investigated. Additionally, animal studies have already provided important insights into the relationship between different stimulation parameters^22–24^, which contribute rich information for hypothesizing which transferable stimulation variants for taVNS studies could enhance the effects on NTS and LC. A double-blind study and stimulation design that allows a flexible change of stimulation location and parameters even during a session under constant stimulation sensation, could provide more concrete data on potential taVNS effects on pupil dilation. Given the susceptibility of pupillometry to the sensations of taVNS, a more direct method could be functional magnetic resonance imaging (fMRI) during taVNS (see Ludwig et al.^1^ for review); however, this could also be affected by attention modulations and different sensations. Furthermore, in the clinical context, it is important to consider changes in the effect of NE release when externally modulating NE release by taVNS (see Ludwig et al.^1^ for review) considering the perception of sensations due to the stimulation. Nonetheless, even not all parameters can be systematically varied simultaneously, and sensations cannot be held perfectly constant for each individual, striving for comparable sham stimulation location, comparable sensations and statistically accounting for sensation variability is an approach that should be pursued more frequently to study the effects of different taVNS stimulation parameters in humans.

## Methods

### Subjects

Twenty-four younger healthy subjects (12 females; 22.96 +/− 2.24 yrs.) were recruited through advertisements via university’s mailing list as well as flyer distributions in Magdeburg. Subjects were included if they were between 20 and 30 years old, German speaking, had a BMI < 27, with low levels of alcohol and cigarette consumption. In addition, subjects were stratified into sporty (more than 3 times a week sport in the last 4 weeks) vs. non-sporty (less than 2 times a week sport in the last 4 weeks) as the whole experiment also included the acquisition of heart-rate variability (HRV) which varies in athletes compared to no athletes^42^. Exclusion criteria included cold symptoms, neurological (stroke, epilepsy, traumatic brain injury, syncope) as well as psychiatric (eating disorder, major depressive disorder, schizophrenia, bipolar disorder, any anxiety disorder, posttraumatic stress disorder) and other disorders (e.g., diabetes, alcohol dependence and/or drug use) as well as heart and eye diseases. Telephone screenings were conducted to verify the eligibility of those interested in the study. Subjects were asked to eat a light, healthy breakfast (no industrial sugar), not to drink caffeine and not to smoke on the day of the experiment, as well as not to drink alcohol on the day of the experiment and the day before.

#### Procedure

The study was conducted as a sham-controlled, single-blind, within-subject, counterbalanced, randomized design using a one-day stimulation protocol. At the beginning of each session subjects underwent a HRV baseline measurement, which was repeated halfway through the whole and at the end of the experiment (see Fig. 2). Subsequently, the subjects were able to try out the taVNS themselves to become familiar with the device and to adjust the highest stimulation intensity (see section *Transcutaneous auricular vagus nerve stimulation*), which was accompanied by a subjective evaluation of the perception using a visual analog scale (VAS) (see section *Visual Analog Scale*). Specifically, it was instructed before, that the stimulation of the ear can be perceived as a harmless tingling in various areas. Additionally, the entire ear was cleaned and not just a specific stimulation area. Furthermore, the repositioning of the electrodes was covered by the story that the cream dries on the electrode after a certain time. This procedure ensured that subjects did not question why the electrodes were being reapplied for real and sham stimulation. The study consisted of two parts: (1) emotional memory task and (2) resting state task. During the performance of the emotional memory task as well as during the presentation of the fixation-cross during the resting state task, subjects received real and sham stimulation while changes in pupil dilation and HRV were recorded in parallel. Immediately after the encoding sessions of the emotional memory test, an early recognition test was performed on the same day, and 24 hours later, a delayed recognition test was performed, both without stimulation (see Fig. 2). Importantly, subjective perceptions of sensations (VAS rating) as well as query of the state of health (potential side effects) (see Supplementary Table S8) were systematically recorded after each stimulation session. The present article focuses only on the changes in pupil dilation due to taVNS during the resting state task.

#### Transcutaneous auricular vagus nerve stimulation (taVNS)

TaVNS was delivered using tVNS Technologies nextGen research device (tVNS R, tVNS Technologies GmbH), which is connected via Bluetooth Low Energy (BLE) connection with an android-based application (BOLZIT, Software development and IT Services) to a) *individually set stimulation parameters* (tVNS Research App) and b) *check the applied stimulation intensity and duration* (tVNS Patient App). The ear electrode “legacy” (tVNS Technologies GmbH) was used, as the size of the electrode holder frame can be adjusted individually. Importantly, the tVNS R device can be connected to a “tVNS Manager” (BOLZIT) console application for Windows 10, which allows time-synchronous stimulation with the required design experiments via an HTTP request. The electrodes were placed on the **left ear** (see Fig. 3): At the **cymba conchae** for **real taVNS**, which seems to be innervated exclusively by the auricular branch of the vagus nerve (ABVN)^43^ and at the **earlobe** for **sham taVNS**, which is not innervated by the ABVN^4,27,43^ and seems to not induce functional activation in the target brain areas, like LC and NTS, following taVNS^2^. For real and sham stimulation, the anode was placed more rostrally. Prior to the electrode placement, the ear was cleaned with *disinfectant alcohol* and afterwards a small amount of EC2+, Grass *electrode conductive cream* (https://www.cnsac-medshop.com/de/ec2-elektrodenleitcreme/) on the electrodes was used to assure optimal conductance. Subsequently, the subjects were able to test the taVNS themselves with a frequency of 25 Hz, a pulse width of 250 μs and a stimulation cycle of 5 s on vs off stimulation. The intensity started at 1 mA and subjects were allowed to go as high as possible at a reasonable pace. At the highest level, subjects rated the subjective intensity on a VAS (see section *Visual Analog Scale*). Low and high **intensities** *(3 mA vs. 5 mA)* as well as **frequencies** *(10 Hz vs. 25 Hz)* were tested systematically within subjects delivered as biphasic square pulses at a **pulse width** of 250 μs during **phasic stimulation** of *3 s ON* and *15 s OFF stimulation*. A priori, it was determined that subjects who did not reach 5 mA as the highest intensity would receive 3 mA as highest and 1.5 mA as lowest intensity, which in the end applied to 7 out of 24 subjects (see Supplementary Fig. S1). Precise control of all BLE-capable devices was important: In the first step, parameters were set using the tVNS Research app, the BLE connection was then removed so that the BLE connection to the taVNS Manager could be guaranteed. Successful stimulation throughout the experiment could be guaranteed as the taVNS Manager sends messages when the stimulation is on and off according to the set duration, which is additionally accompanied by a continuous light of the tVNS R device while the stimulation is on.

**Figure 3.**
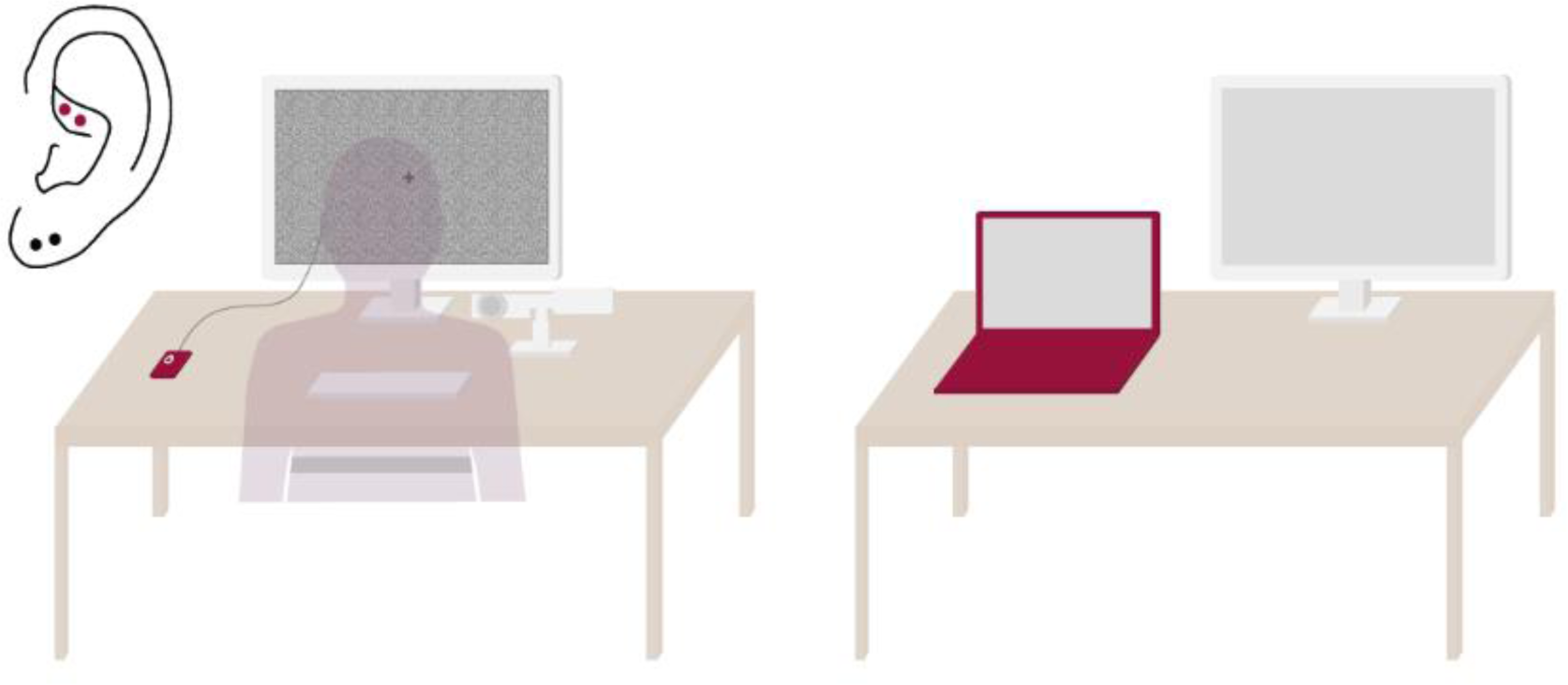
Experimental set-up of the resting-state task. The experimental set-up enabled a time-synchronous phasic stimulation during the resting state task while changes in heart-rate variability (HRV; belt, left) and pupil dilation (eyetracker camera, left) were recorded in parallel on one computer (red laptop, right). This computer was connected to an extended screen (left) on which the task was presented. Additionally, the computer received stimulation inputs via an HTTP request that turned on or off the taVNS R stimulator (red square, left) in sections programmed for the task. In addition, a second screen (right) allowed to control of pupil recordings. The electrodes were placed on the **left** ear: At the **cymba conchae** for **real taVNS** (red dots) and at the **earlobe** for **sham taVNS** (black dots). For all subjects, the same constant ambient light continued to be applied throughout the whole experiment and background’s brightness variations were controlled with a grayish background image to prevent interference of luminance changes with pupillometric recordings.

#### Resting state task

The resting state task consisted of two blocks (randomisation and counterbalancing of real and sham stimulation between subjects) of four sessions, each with 60 trials. The subjects were instructed to focus their gaze on a grey fixation cross throughout the task (see Fig. 3). Each trial began with a grey fixation cross during which stimulation was turned on for 3 s, followed by another grey fixation cross during which stimulation was turned off for 15 s on average (13 – 17 s). To prevent interference with pupillometric recordings, the background’s brightness variations were controlled with a greyish background image^18^ (see Fig. 2). Both high and low frequency and intensity were tested in the 4 sessions randomized within and between subjects for real and sham stimulations. Between the single sessions, parameters were adjusted, and subjects were able to take a break (5-10 min). Those who needed the longer break were asked to walk around in the hallway outside the lab room to ensure sufficient attentional focus for the next session. Between the two blocks there was a break of 20 min, if necessary, up to 30 min. The total task duration was 2 blocks * (18 min + 10 min post HRV measurement * 4 conditions) 4h 10 min, with the stimulation lasting a total of 12 min (60 trials * 4 conditions * 3 s) during each block (3 min per condition). The experiment was controlled by custom MATLAB code (Math Works, www.mathworks.com) using Psychtoolbox 3 (www.psychtoolbox.org), while stimulation could be controlled in a time-synchronized manner with the experiment via the “tVNS Manager”. Thus, messages were forwarded via the “tVNS Manager”, which were integrated within the MATLAB code, so that the stimulation was either switched on or off per trial within a loop.

### Visual Analog Scale (VAS)

Subjects were asked to rate how pleasant or unpleasant each stimulation session was perceived after stimulation, based on a visual analog scale (VAS)^44^ ranging from (1) very pleasant to (10) very unpleasant. Since it has been shown that the perception of sensations differs between real and sham stimulation, the VAS is often kept constant as a controlling factor in many studies and the individual intensity is allowed to vary for each subject based on e.g., a “tingling” sensation below the pain threshold^25,45,46^. However, because we systematically tested a fixed set of different intensities and frequencies, we could not hold VAS constant but could document effects of the different parameters on perception of sensations (see Supplementary Fig. S3).

### State of health

The state of health queried in each case after stimulation to control for potential side effects are shown in Supplementary Table S8. The following items were asked: (1) headache, (2) nausea, (3) tiredness, (4) dizziness, (5) tingling sensation at the previously stimulated area, (6) feeling of heat at the previously stimulated area, (7) reddening of the skin at the previously stimulated area, (8) skin irritation at the previously stimulated site, (9) impaired concentration, (10) itching at the previously stimulated area. Subjects indicated on a 4-point scale (0: not at all – 3: strong) to what extent they perceived potential side effects. The reported sensations did not differ between real (M = 0.21, SD = 0.13) and sham (M = 0.17, SD = 0.10) stimulation (F(1,9) = 3.30, p = 0.10) and between low (M = 0.18, SD = 0.12) and high (M = 0.18, SD = 0.21) frequency (F(1,9) = 0.13, p = 0.72). There was a significant difference between low (M = 0.16, SD = 0.11) and high (M = 0.20, SD = 0.12) intensity (F(1,9) = 5.34, p = 0.05). Overall, it can be concluded that there were no side effects due to the stimulation and that the minimal impairments were rather due to the long measurement day and the monotonous resting state task (e.g., item tiredness (3) and concentration (9)) than to the stimulation itself. Thus, the stimulation can be considered safe, which is in line with previous reports^25^.

#### Pupil data acquisition

Changes in pupil diameter were continuously recorded monocularly from the left eye at a sampling rate of 1000 Hz using a desked-based infrared EyeLink 1000 eyetracker (SR Research, www.sr-research.com) with a chin rest. The centroid measure of pupil change was chosen to provide more accurate estimates of changes in pupil dilation over time. The recording of pupillometry was controlled by custom-made scripts in MATLAB 2020b (Math Works, www.mathworks.com) using Psychtoolbox 3 (www.psychtoolbox.org) and the Eyelink add-in toolbox for eyetracker control. For all subjects, the same constant ambient light continued to be applied throughout the whole experiment. At the start of the experiment the camera was calibrated using 5-point calibration.

#### Pupil data analysis

Pupil data were pre-processed and analysed using custom-made scripts in MATLAB 2020b (Math Works, www.mathworks.com). For pre-processing, pupil data were segmented 200 ms before and 17.5 s after trial onset. To clean pupil data from artefacts and blinks, the data was further processed following recommendations in Mathot^47^. First, the signal was smoothed using a moving Hanning window (15 ms) average. A velocity profile was then created based on the smoothed signal to detect, using a threshold of mean-standard deviation, to identify the beginning (velocity is below a threshold) and the end of a blink (velocity is above a threshold) as well as closed eyes (velocity is zero). Since the blink period can be underestimated^47^ 40 ms were additionally subtracted from the beginning time and added to the end time. All defined artefacts and blinks were set to NaN, summarised and then linearly interpolated. For the analyses, only trials whose raw signal was 70% free of blinks and artefacts, allowing 30% for interpolated data were included. Variations in trial numbers per condition were observed following artifact correction (real stimulation: M = 53.8, SD = 10.9; sham stimulation: M = 58.5, SD = 5.23; see Supplementary Results 1). Finally, all trials were also verified by visual inspection. Pupil data were baseline-corrected (200 ms before stimulation onset) as well as individually z-scored to allow comparison of task conditions independent of individual differences in pupil dilation size^48,49^. The z standardised and baseline corrected data were analysed separately in three-time windows (see Fig. 1): **(I)** the 3 s during on stimulation (**‘on stimulation’**), **(II)** the first 3 s during off stimulation (**‘immediate response’**) and **(III)** the subsequent last 10 s during off stimulation (**‘delayed response’**).The selection of the three different time windows was based on 3 s of phasic stimulation as well as an also equal length of an immediate response followed by a longer delayed response due to the trial duration.

#### Statistical analysis

Statistical analyses were conducted in R version 4.2.2 (R Core Team, 2022) using RStudio version (RStudio Team, 2022) and graphs were created using the package ggplot2^50^. The mean value of the respective items for potential side effects (state of health) as well as the perception of sensations (VAS rating) for (I) ‘on stimulation’ were analysed across all subjects by using aov_result() function for repeated-measures ANOVA ({afex} package^49^) and lsmeans() function ({emmeans} package^56^). Additionally, Pearson correlation coefficients between VAS and pupil dilation (averaged per subject across trials) during (**I**) ‘on stimulation’ were calculated by using cor.test() and corrected for outliers based on interquartile range (1.5*IQR).

Furthermore, changes in pupil dilation were analysed based on a fitted linear mixed-effects (LMM) model by using the {lme4} package^51^, following a forward model selection approach. Thereby, a distinct model was fitted for each time window (**I-III**) (see Supplementary Table S1) using the same dummy coded variables (see below). LMM allows to account for the nested structure of the repeated measured data and for using a random intercept for each subject to account for interindividual differences in mean pupil responses. Additionally, this approach allows modelling the data at the level of individual trials to account for time-on-task effects on pupil dilations.

The model comparisons were conducted using the anova() function ({lme4} package^51^) with likelihood-ratio chi-squared tests. AIC (Akaike Information Criterion) values of the best model for statistical modelling and model selection were reported. In general, models with lower AIC values are indicative of a superior trade-off between data explanation and prevention of overfitting, in comparison to alternative assessed models^52^. To assess the relevant assumptions of LMM, check_model() function ({performance} package^53^) was used to investigate linearity, homogeneity of variance, influential observations, collinearity, normality of residuals and of random effects (https://osf.io/va64p/). The significance of predictors on the goodness of fit of the model was assessed using Anova() function ({car} package^54^), which computes type-II analysis-of-variance tables for mixed-effects models and provides likelihood-ratio Chi-Square statistics. The significance of the deviance of individual groups from the intercept was assessed using summary() function ({lmerTest} package^55^), which calculates model’s coefficients, standard errors, t-values, and p-values associated with each coefficient. In addition, to incorporating a random intercept ‘ID’, which accounts for interindividual variations in mean pupil change, and the inclusion of the trial number variable ‘trials’ to capture the impact of ‘time-on-task’ on pupil dilations, the forward model selection approach initially considered variables related to the distinction between real and sham ***‘stimulation’*** *[real (1) vs. sham (0)]*, as well as differences in ***‘frequency’*** *[high (1) vs. low (0)]* and ***‘intensity’*** *[high (1) vs. low (0)]* (see Supplementary Table S1). It was furthermore investigated whether incorporating interactions between stimulation, frequency and intensity further improved the model (see Supplementary Table S1-S4*; anova(m0, m1, m2, m3, m4, m5, m6, m7)*). This did not lead to a significant improvement in the model fit for the time windows (I-III). Hence, it was determined that the best model from the first step (see results) for each distinct model at every given time (**I-III**) point was **model m_4**:

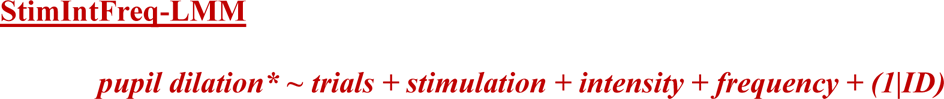

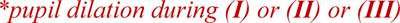

Second, based on **model m_4** the following factors were added stepwise: ***‘VAS’*** ratings as a measure of subjective perception of sensations due to stimulation, ***‘sensitivity’*** *[sensitive (1) vs. not sensitive (0)]* differentiating whether subjects received 3 and 5 mA or 1.5 and 3 mA (see Supplementary Fig. S1), whether subjects received real stimulation first ***‘real_first’*** *[counterbalanced: real (1) before sham (0) stimulation],* in which order the four stimulation combinations were applied **‘*position’*** *[randomised: low mA & low Hz (1), high mA & low Hz (2), low mA & high Hz (3), high mA & high Hz (4)], **gender** [female (1) vs. male (0)* and ***sporty*** *[sporty (1) vs. non-sport (0)].* Subsequently model comparisons were conducted again based on all models without interactions (see results; *anova(m0, m1, m2, m3, m4, m4_1, m4_2, m4_3, m4_4, m4_5, m4_6*)). The best fitting model from the second step for each time point was **model m_4_1**:

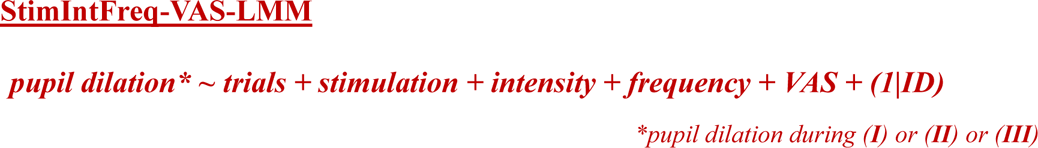

Additionally, random effects for stimulation, frequency and intensity were added stepwise to the model ‘StimIntFreq-VAS-LMM’, while only the model with intensity as a random effect led to evaluable results (see Supplementary Results 2), possibly due to intensity variations yielding strongest stimulation effects.

Third, an exploratory analysis was added to better explain the potential influence of sensory perception (VAS) due to stimulation on pupil dilation controlled for sensitivity:

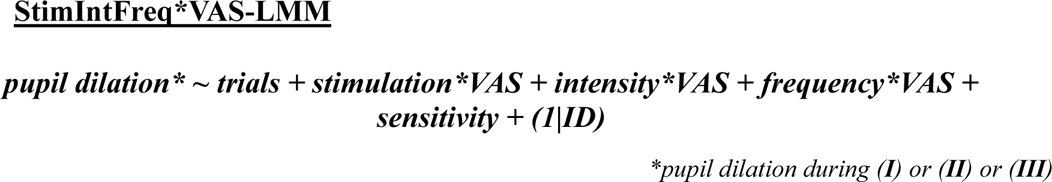

Moreover, emmip() and emtrends() functions ({emmeans} package^56^) were utilized to analyse interaction effects, such as VAS changes along its range with respect to categorical variables (stimulation, frequency, intensity). The function emmip() generates an interaction plot to see how the categorical variable affects the variable over its entire range. The function emtrends() calculates estimated marginal means for different levels of the categorical variable. Distinct models based on the average pupil dilation per session are also reported for each time window (**I-III**) using the same criteria, but without the covariate ‘trial number’ (see Supplementary Table S5-6).

#### Ethics approval, written informed consent, and compensation

The study was approved by the Ethics Committee of medical faculty at the Otto von Guericke University of Magdeburg (reference no. 107/20) and was carried out in accordance with the ethical standards of Helsinki. A written informed consent was obtained from each subject before participation, and they were reimbursed with 90 Euros.

## Acknowledgements

M.L is supported by the federal state of Saxony-Anhalt and the European Regional Development Fund (ERDF) in the Center for Behavioral Brain Sciences (CBBS, ZS/2016/04/78113).

E.D has received financial support for his institution by Deutsche Forschungsgemeinschaft (DFG, German Research Foundation) – ProjectID 425899994 – Sonderforschungsbereiche 1436 (SFB 1436), Human Brain Project, Specific Grant Agreement 3 (SGA3), Deutsche Forschungsgemeinschaft (DFG, German Research Foundation) – Sonderforschungsbereiche 1315 (SFB 1315).

M.J.B is supported by the Deutsche Forschungsgemeinschaft (DFG, German Research Foundation) – Project-ID 425899994 – Sonderforschungsbereiche 1436 (SFB 1436), Center for Behavioral Brain Sciences (CBBS) NeuroNetzwerk 17, and by the German Federal Ministry of Education and Research (BMBF, funding code 01ED2102B) under the aegis of the EU Joint Programme – Neurodegenerative Disease Research (JPND).

D.H is supported by Sonderforschungsbereich 1315, Project B06, Sonderforschungsbereich 1436, Project A08, ARUK SRF2018B-004, CBBS Neural Network (CBBS, ZS/2016/04/78113), and NIH R01MH126971.

## Author Contribution

M.L., D.H. and M.J.B. contributed to the conceptualization and methodology of the study. M.L. carried out the measurements, analysed and visualised the data and wrote the manuscript. C.P. supported with the technical construction and implementation of the taVNS set-up. M.K. advised on the statistical analyses. D.H. and M.J.B. supervised the investigation, analyses, and writing of the original draft. All authors reviewed the manuscript.

## Additional Information Competing Interest

E.D has received payments for his role and works as consultant for Roche, Biogen, RoxHealth and expert testimony for UCL Consultancy, served at scientific advisory boards for EdoN Initiative and Ebsen Alzheimers Center (no payment) and Roche (personal financial support), and is a co-founder of the digital health start-up Neotiv. M.L., C.P., M.K., M.J.B., D.H. have nothing to disclose.

## Data availability

The dataset generated and analysed during the current study is available online at https://osf.io/va64p/ (OpenScience Framework).

## References

1. Ludwig, M., Wienke, C., Betts, M. J., Zaehle, T. & Hämmerer, D. Current challenges in reliably targeting the noradrenergic locus coeruleus using transcutaneous auricular vagus nerve stimulation (taVNS). Autonomic Neuroscience 102900 (2021) doi:10.1016/j.autneu.2021.102900.

2. Yakunina, N., Kim, S. S. & Nam, E.-C. Optimization of Transcutaneous Vagus Nerve Stimulation Using Functional MRI: TRANSCUTANEOUS VNS OPTIMIZATION USING fMRI. Neuromodulation: Technology at the Neural Interface 20, 290–300 (2017).

3. Sharon, O., Fahoum, F. & Nir, Y. Transcutaneous Vagus Nerve Stimulation in Humans Induces Pupil Dilation and Attenuates Alpha Oscillations. J. Neurosci. 41, 320–330 (2021).

4. Butt, M. F., Albusoda, A., Farmer, A. D. & Aziz, Q. The anatomical basis for transcutaneous auricular vagus nerve stimulation. J. Anat. joa.13122 (2019) doi:10.1111/joa.13122.

5. Ruffoli, R. et al. The chemical neuroanatomy of vagus nerve stimulation. Journal of Chemical Neuroanatomy 42, 288–296 (2011).

6. Aston-Jones, G. & Bloom, F. E. Activity of norepinephrine-containing locus coeruleus neurons in behaving rats anticipates fluctuations in the sleep-waking cycle. J. Neurosci. 1, 876–886 (1981).

7. Aston-Jones, G. & Cohen, J. D. AN INTEGRATIVE THEORY OF LOCUS COERULEUS-NOREPINEPHRINE FUNCTION: Adaptive Gain and Optimal Performance. Annual Review of Neuroscience 28, 403–450 (2005).

8. Berridge, C. W. & Waterhouse, B. D. Review. (2003).

9. Clayton, E. C., Rajkowski, J., Cohen, J. D. & Aston-Jones, G. Phasic activation of monkey locus ceruleus neurons by simple decisions in a forced-choice task. J Neurosci 24, 9914–9920 (2004).

10. Florin-Lechner, S. M., Druhan, J. P., Aston-Jones, G. & Valentino, R. J. Enhanced norepinephrine release in prefrontal cortex with burst stimulation of the locus coeruleus. Brain Research 742, 89–97 (1996).

11. Joshi, S. & Gold, J. I. Pupil size as a window on neural substrates of cognition. Trends Cogn Sci 24, 466–480 (2020).

12. Joshi, S., Li, Y., Kalwani, R. M. & Gold, J. I. Relationships between Pupil Diameter and Neuronal Activity in the Locus Coeruleus, Colliculi, and Cingulate Cortex. Neuron 89, 221–234 (2016).

13. Reimer, J. et al. Pupil fluctuations track rapid changes in adrenergic and cholinergic activity in cortex. Nat Commun 7, 13289 (2016).

14. Murphy, P. R., O’Connell, R. G., O’Sullivan, M., Robertson, I. H. & Balsters, J. H. Pupil diameter covaries with BOLD activity in human locus coeruleus. Human Brain Mapping 35, 4140–4154 (2014).

15. Varazzani, C., San-Galli, A., Gilardeau, S. & Bouret, S. Noradrenaline and dopamine neurons in the reward/effort trade-off: a direct electrophysiological comparison in behaving monkeys. J Neurosci 35, 7866–7877 (2015).

16. de Gee, J. W. et al. Dynamic modulation of decision biases by brainstem arousal systems. eLife 6, e23232 (2017).

17. Breton-Provencher, V. & Sur, M. Active control of arousal by a locus coeruleus GABAergic circuit. Nat Neurosci 22, 218–228 (2019).

18. Mathôt, S., Fabius, J., Van Heusden, E. & Van der Stigchel, S. Safe and sensible preprocessing and baseline correction of pupil-size data. Behav Res 50, 94–106 (2018).

19. Hall, C. A. & Chilcott, R. P. Eyeing up the Future of the Pupillary Light Reflex in Neurodiagnostics. Diagnostics 8, 19 (2018).

20. Samuels, E. & Szabadi, E. Functional Neuroanatomy of the Noradrenergic Locus Coeruleus: Its Roles in the Regulation of Arousal and Autonomic Function Part I: Principles of Functional Organisation. CN 6, 235–253 (2008).

21. Manta, S., El Mansari, M., Debonnel, G. & Blier, P. Electrophysiological and neurochemical effects of long-term vagus nerve stimulation on the rat monoaminergic systems. International Journal of Neuropsychopharmacology 16, 459–470 (2013).

22. Mridha, Z. et al. Graded recruitment of pupil-linked neuromodulation by parametric stimulation of the vagus nerve. Nat Commun 12, 1539 (2021).

23. Collins, L., Boddington, L., Steffan, P. J. & McCormick, D. Vagus nerve stimulation induces widespread cortical and behavioral activation. Current Biology 31, 2088–2098.e3 (2021).

24. Hulsey, D. R. et al. Parametric characterization of neural activity in the locus coeruleus in response to vagus nerve stimulation. Experimental Neurology 289, 21–30 (2017).

25. Farmer, A. D. et al. International Consensus Based Review and Recommendations for Minimum Reporting Standards in Research on Transcutaneous Vagus Nerve Stimulation (Version 2020). Front. Hum. Neurosci. 14, 568051 (2021).

26. Lloyd, B., Wurm, F., de Kleijn, R. & Nieuwenhuis, S. Short-term transcutaneous vagus nerve stimulation increases pupil size but does not affect EEG alpha power: a replication. http://biorxiv.org/lookup/doi/10.1101/2023.03.08.531479 (2023) doi:10.1101/2023.03.08.531479.

27. Burger, A. M., D’Agostini, M., Verkuil, B. & Van Diest, I. Moving beyond belief: A narrative review of potential biomarkers for transcutaneous vagus nerve stimulation. Psychophysiology 57, e13571 (2020).

28. D’Agostini, M. et al. Short bursts of transcutaneous auricular vagus nerve stimulation enhance evoked pupil dilation as a function of stimulation parameters. Cortex 159, 233–253 (2023).

29. Sclocco, R. Stimulus frequency modulates brainstem response to respiratory-gated transcutaneous auricular vagus nerve stimulation. Brain Stimulation 9 (2020).

30. Urbin, M. A. et al. Electrical stimulation of the external ear acutely activates noradrenergic mechanisms in humans. Brain Stimulation 14, 990–1001 (2021).

31. Loerwald, K. W., Borland, M. S., Rennaker, R. L., Hays, S. A. & Kilgard, M. P. The interaction of pulse width and current intensity on the extent of cortical plasticity evoked by vagus nerve stimulation. Brain Stimulation 11, 271–277 (2018).

32. Kahneman, D. Attention and effort. (Prentice-Hall, Inc, 1973).

33. Miller, A. L., Gross, M. P. & Unsworth, N. Individual differences in working memory capacity and long-term memory: The influence of intensity of attention to items at encoding as measured by pupil dilation. Journal of Memory and Language 104, 25–42 (2019).

34. Sara, S. J. Noradrenergic Modulation of Selective Attention: Its Role in Memory Retrieval. Annals of the New York Academy of Sciences 444, 178–193 (1985).

35. Aston-Jones, G., Rajkowski, J., Kubiak, P. & Alexinsky, T. Locus coeruleus neurons in monkey are selectively activated by attended cues in a vigilance task. J Neurosci 14, 4467– 4480 (1994).

36. Evans, M. S., Verma-Ahuja, S., Naritoku, D. K. & Espinosa, J. A. Intraoperative human vagus nerve compound action potentials. Acta Neurol Scand 110, 232–238 (2004).

37. Chang, Y.-C. et al. Quantitative estimation of nerve fiber engagement by vagus nerve stimulation using physiological markers. Brain Stimulation 13, 1617–1630 (2020).

38. Safi, S., Ellrich, J. & Neuhuber, W. Myelinated Axons in the Auricular Branch of the Human Vagus Nerve: Auricular Vagus Nerve Branch. Anat. Rec. 299, 1184–1191 (2016).

39. Krahl, S. E., Senanayake, S. S. & Handforth, A. Destruction of Peripheral C-Fibers Does Not Alter Subsequent Vagus Nerve Stimulation-Induced Seizure Suppression in Rats. Epilepsia 42, 586–589 (2001).

40. Brodin, P. DIfferential inhibition of A, B and C fibres in the rat vagus nerve by lidocaine, eugenol and formaldehyde. Archives of Oral Biology 30, 477–480 (1985).

41. Zanos, T. P. et al. Identification of cytokine-specific sensory neural signals by decoding murine vagus nerve activity. Proc. Natl. Acad. Sci. U.S.A. 115, (2018).

42. Kiss, O. et al. Detailed heart rate variability analysis in athletes. Clin Auton Res 26, 245–252 (2016).

43. Peuker, E. T. & Filler, T. J. The nerve supply of the human auricle. Clin. Anat. 15, 35– 37 (2002).

44. Yeung, A. W. K. & Wong, N. S. M. The Historical Roots of Visual Analog Scale in Psychology as Revealed by Reference Publication Year Spectroscopy. Front Hum Neurosci 13, 86 (2019).

45. Ferstl, M. et al. Non-invasive vagus nerve stimulation boosts mood recovery after effort exertion. 26.

46. Müller, F. K., Teckentrup, V., Kühnel, A., Ferstl, M. & Kroemer, N. B. Acute vagus nerve stimulation does not affect liking or wanting ratings of food in healthy participants. http://biorxiv.org/lookup/doi/10.1101/2021.03.26.437062 (2021) doi:10.1101/2021.03.26.437062.

47. Mathôt, S. A simple way to reconstruct pupil size during eye blinks. 4.

48. Hämmerer, D. et al. Emotional arousal and recognition memory are differentially reflected in pupil diameter responses during emotional memory for negative events in younger and older adults. Neurobiology of aging 58, 129–139 (2017).

49. Hämmerer, D. et al. Older adults fail to form stable task representations during model-based reversal inference. Neurobiology of Aging 74, 90–100 (2019).

50. Wickham, H., et al. ggplot2: Create Elegant Data Visualisations Using the Grammar of Graphics. (2023).

51. Bates, D., Mächler, M., Bolker, B. & Walker, S. Fitting Linear Mixed-Effects Models Using lme4. J. Stat. Soft. 67, (2015).

52. Vrieze, S. I. Model selection and psychological theory: A discussion of the differences between the Akaike Information Criterion (AIC) and the Bayesian Information Criterion (BIC). Psychol Methods 17, 228–243 (2012).

53. Lüdecke, D., Ben-Shachar, M., Patil, I., Waggoner, P. & Makowski, D. performance: An R Package for Assessment, Comparison and Testing of Statistical Models. JOSS 6, 3139 (2021).

54. Fox, J., et al. car: Companion to Applied Regression. (2023).

55. Kuznetsova, A., Brockhoff, P. B. & Christensen, R. H. B. lmerTest Package: Tests in Linear Mixed Effects Models. J. Stat. Soft. 82, (2017).

56. Lenth, R. V. R package emmeans: Estimated marginal means. (2023).

