## supplemental Files for "Evaluating phasic transcutaneous vagus nerve stimulation (taVNS) with pupil dilation: the importance of stimulation intensity and sensory perception"

[a]Institute of Cognitive Neurology and Dementia Research, Otto-von-Guericke University Magdeburg, Magdeburg, Germany; [b]CBBS Center for Behavioral Brain Sciences, Magdeburg, Germany; [c]Otto-von-Guericke University Magdeburg, Magdeburg, Germany; [d]Institute for Neuromodulation and Neurotechnology, University Hospital and University of Tuebingen, Tuebingen, Germany; [e]German Center for Neurodegenerative Diseases (DZNE), Otto-von-Guericke University Magdeburg, Magdeburg, Germany;[f]Institute of Cognitive Neuroscience, University College London, London, UK; [g]The Wellcome Trust Centre for Neuroimaging, University College London, London, UK; [h]Department of Psychology, University of Innsbruck

### Methods

#### Subjects

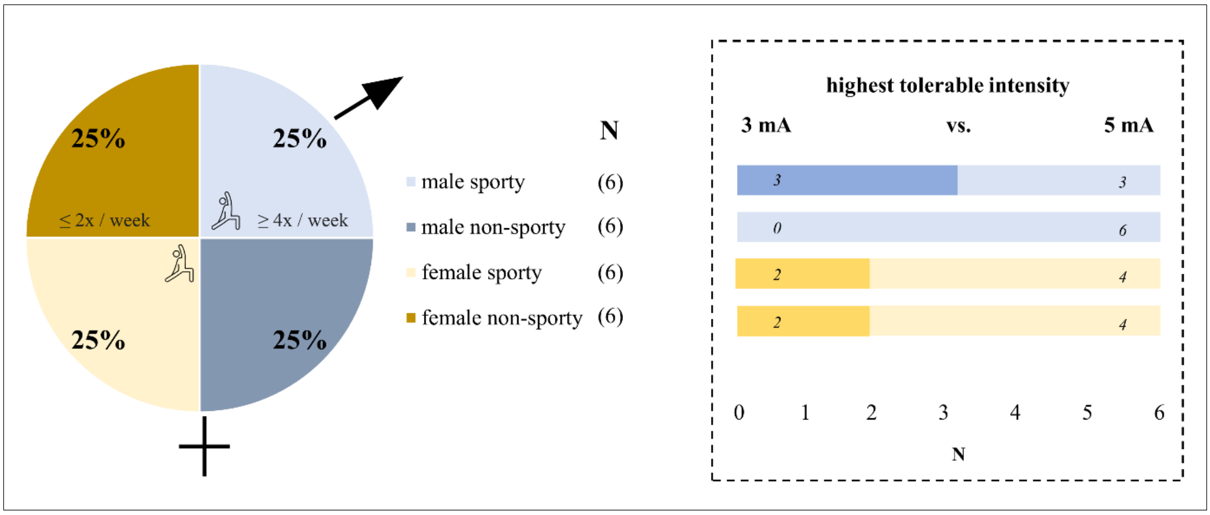


**Supplementary Figure S1**. Twenty-four subjects participated in the study, of which 50% female (ochre) and 50% male (blue) and of which 25% sporty vs. 25% non-sporty respectively, based on a prior established criterion (sporty = more than 3 times a week sport in the last 4 weeks; non-sporty = less than 2 times a week sport in the last 4 weeks). It was predetermined that subjects would receive 3 mA as the highest and 1.5 mA as the lowest intensity if they did not reach 5 mA as the highest intensity, which was the case for 7 of the 24 subjects.

**Determination of sample size**. A study with sixteen healthy younger adults were able to show combined behavioural and electrophysiological evidence for GABAergic neuromodulation by taVNS with an effect size of $\eta 2 = 0.271$; $F(1,15) = 5.57$, $p = .032$ (Keute et al., 2018). Based on our a-prior calculation in G*Power to determine the number of subjects, we used a medium effect size (0.25, based on Cohen´s guidelines) and set a conventional alpha level of 0.05 and power of 0.80 to ensure a high probability of detecting true effects, leading to a sample size of N = 24 subjects.

### Supplementary Results 1

#### Pupil data analysis

**Excluded trials.** Variations in trial numbers per condition were observed following artifact correction. Specifically, sham stimulation (M = 58.5, SD = 1.07) had more trials compared to real stimulation (M = 53.8, SD = 1.91), $F(1,23) = 15.89$, $p < 0.001$; high frequency stimulation (M = 57.8, SD = 1.27) had more trials than low frequency stimulation (M = 54.5, SD = 1.65), $F(1,23) = 24.33$, $p < 0.001$ and high intensity stimulation (M = 57.2, SD = 1.32) had more trials than low intensity stimulation (M = 55.0, SD = 1.56), $F(1,23) = 29.98$, $p < 0.001$. Likewise, there was a significant interaction between stimulation condition and frequency ($F(1,23) = 24.87$, $p < 0.001$) (*high frequency during real stimulation (M = 56.8, SD = 1.63) vs. low* *frequency during real stimulation (M = 50.8, SD = 2.31), t(23) = 5.15, p = 0.0002; low frequency during real stimulation (M = 50.8, SD = 2.31) vs. low frequency during sham stimulation (M = 58.2, SD = 1.15), t(23) = -4.74, p = 0.0005).* There was also a significant interaction between stimulation condition and intensity (F (1,23) = 31.13, p < 0.001) (*high intensity during real stimulation (M = 55.9, SD = 1.67) vs. low intensity during real stimulation (M = 51.6, SD = 2.19), t(23) = 5.54, p = 0.0001; low intensity during real stimulation (M = 51.6, SD = 2.19) vs. low intensity during sham stimulation (M = 58.4, SD = 1.08), t(23) = -4.5, p = 0.0009).* Moreover there was a significant interaction between intensity and frequency (F (1,23) = 30, p < 0.001) *(low intensity and high frequency (M = 57.8, SD = 1.27) vs. low intensity low frequency (M = 52.3, SD = 1.94), t(23) = 5.19, p = 0.0002; high intensity and low frequency (M = 56.6, SD = 1.40) vs. low intensity and low frequency (M = 52.3, SD = 1.94), t(23) = 5.48, p = 0.0001).* Finally, there was a significant three-way-interaction between stimulation condition, frequency and intensity (F (1,23) = 31.13, p < 0.001).

**Statistical analysis.**

**Distinct models for each time window (I-III)**

| **Models**: |
|---|
| m0: Pupil ~ (1 \| ID) |
| m1: Pupil ~ trials + (1\|ID) |
| m2: Pupil ~ trials + stimulation + (1\|ID) |
| m3: Pupil ~ trials + stimulation + freq + (1\|ID) |
| m4: Pupil ~ trials + stimulation + freq + int + (1\|ID) |
| m5: Pupil ~ trials + stimulation *freq + int + (1\|ID) |
| m6: Pupil ~ trials + stimulation *freq + stimulation *int + (1\|ID) |
| m7: Pupil ~ trials + stimulation *freq + stimulation *int + int*freq + (1\|ID) |
| m4_1: Pupil ~ trials + stimulation + freq + int + vas + (1\|ID) |
| m4_2: Pupil ~ trials + stimulation + freq + int + vas + sensitive + (1\|ID) |
| m4_3: Pupil ~ trials + stimulation + freq + int + vas + sensitive + real_first + (1\|ID) |
| m4_4: Pupil ~ trials + stimulation + freq + int + vas + sensitive + real_first + position + (1\|ID) |
| m4_5: Pupil ~ trials + stimulation + freq + int + vas + sensitive + real_first + position + gender + (1\|ID) |
| m4_6: Pupil ~ trials + stimulation + freq + int + vas + sensitive + real_first + position + gender + sporty + (1\|ID) |

**Supplementary Table S1.** Changes in pupil dilation were analysed based on a fitted linear mixed-effects (LMM) model by using the {lme4} package (Bates et al., 2015), following a forward model selection approach. Thereby, a distinct model was fitted for each time window (**I-III**) using the same dummy coded variables *(**stimulation** [real (1) vs. sham (0)], **frequency** [high (1) vs. low (0)] (freq), **intensity** [high (1) vs. low (0)](int), **trial number** (trials), **sensitivity** [sensitive (1) vs. not sensitive (0)], **real_first** [counterbalanced: real (1) before sham (0) stimulation], **position** (four different stimulation combination possibilities), **sporty** [sporty (1) vs. non-sport (0)], **gender** [female (1) vs. male (0)] and **VAS).***

**Comparison of multiple statistical models for (I) – 3 sec during on stimulation**

| | npar | AIC | BIC | logLik | deviance | Chisq | Df | Pr(>Chisq) |
|---|---|---|---|---|---|---|---|---|
| **m0** | 3 | 29526 | 29548 | -14760 | 29520 | | | |
| **m1** | 4 | 29527 | 29556 | -14759 | 29519 | 1.28 | 1 | 0.26 |
| **m2** | 5 | 29512 | 29549 | -14751 | 29502 | 16.42 | 1 | 5.09e-05 *** |
| **m3** | 6 | 29505 | 29548 | -14746 | 29493 | 9.62 | 1 | 0.0019 ** |
| **m4** | **7** | **29404** | **29455** | **-14695** | **29390** | **102.95** | **1** | **<2.2e-16 *** ** |
| **m4_1** | **8** | **29367** | **29425** | **-14675** | **29351** | **38.76** | **1** | **4.79e-10 *** ** |
| **m5** | 8 | 29405 | 29464 | -14695 | 29389 | 0 | 0 | |
| **m6** | 9 | 29407 | 29473 | -14694 | 29389 | 0.39 | 1 | 0.53 |
| **m4_2** | 9 | 29369 | 29434 | -14675 | 29351 | 38.31 | 0 | |
| **m7** | 10 | 29408 | 29481 | -14694 | 29388 | 0 | 1 | 1 |
| **m4_3** | 10 | 29370 | 29443 | -14675 | 29350 | 38.42 | 0 | |
| **m4_4** | 13 | 29373 | 29468 | -14674 | 29347 | 2.64 | 3 | 0.45 |
| **m4_5** | 14 | 29375 | 29477 | -14674 | 29347 | 0.06 | 1 | 0.81 |
| **m4_6** | 15 | 29377 | 29486 | -14674 | 29347 | 0.04 | 1 | 0.84 |
| Signif. codes: 0 '***' 0.001 '**' 0.01 '*' 0.05 '.' 0.1 ' ' 1 | | | | | | | | |

**Supplementary Table S2**. Comparison of multiple statistical models for time window **(I) – 3 sec during on stimulation**. The overview shows *npar*, which represents the number of parameters in each model, *AIC* (Akaike Information Criterion), *BIC* (Bayesian Information Criterion), *logLik* (Log-Likelihood), *deviance* which was used to assess the goodness of fit of a model, *Chisq* (Chi-Square),which represents the difference in deviance between the current model and the previous one, *Df* (Degrees of Freedom), which represents the degrees of freedom associated with the Chi-Square test and *Pr(>Chisq)* which represents p-values associated with the Chi-Square test. Model m4 (red) was the best fitting model and m4_7 (red) was the second-best fitting model.

**Comparison of multiple statistical models for (II) – immediate response**

| | npar | AIC | BIC | logLik | deviance | Chisq | Df | Pr(>Chisq) |
|---|---|---|---|---|---|---|---|---|
| **m0** | 3 | 37773 | 37795 | -18884 | 37767 | | | |
| **m1** | 4 | 37775 | 37804 | -18884 | 37767 | 0.2370 | 1 | 0.6263508 |
| **m2** | 5 | 37763 | 37799 | -18876 | 37753 | 142.569 | 1 | 0.0002*** |
| **m3** | 6 | 37762 | 37806 | -18875 | 37750 | 23.924 | 1 | 0.1219234 |
| **m4** | **7** | **37688** | **37739** | **-18837** | **37674** | **765.394** | **1** | **< 2.2e-16*** ** |
| **m4_1** | **8** | **37659** | **37717** | **-18822** | **37643** | **308.109** | **1** | **2.84e-08*** ** |
| **m5** | 8 | 37688 | 37746 | -18836 | 37672 | 0.0000 | 0 | |
| **m6** | 9 | 37687 | 37752 | -18834 | 37669 | 31.968 | 1 | 0.074 |
| **m4_2** | 9 | 37661 | 37727 | -18822 | 37643 | 255.596 | 0 | |
| **m7** | 10 | 37688 | 37761 | -18834 | 37668 | 0.0000 | 1 | 1 |
| **m4_3** | 10 | 37662 | 37735 | -18821 | 37642 | 259.133 | 0 | |
| **m4_4** | 13 | 37660 | 37755 | -18817 | 37634 | 84.556 | 3 | 0.04 |
| **m4_5** | 14 | 37662 | 37764 | -18817 | 37634 | 0.0560 | 1 | 0.81 |
| **m4_6** | 15 | 37664 | 37773 | -18817 | 37634 | 0.3817 | 1 | 0.54 |
| Signif. codes: 0 '***' 0.001 '**' 0.01 '*' 0.05 '.' 0.1 ' ' 1 | | | | | | | | |

**Supplementary Table S3**. Comparison of multiple statistical models for the **(II) – immediate response**. The overview shows *npar*, which represents the number of parameters in each model, *AIC* (Akaike Information Criterion), *BIC* (Bayesian Information Criterion), *logLik* (Log-Likelihood), *deviance* which was used to assess the goodness of fit of a model, *Chisq* (Chi-Square),which represents the difference in deviance between the current model and the previous one, *Df* (Degrees of Freedom), which represents the degrees of freedom associated with the Chi-Square test and *Pr(>Chisq)* which represents p-values associated with the Chi-Square test. Model m4 (red) was the best fitting model and m4_7 (red) was the second-best fitting model.

**Comparison of multiple statistical models for (III) – delayed response**

| | npar | AIC | BIC | logLik | deviance | Chisq | Df | Pr(>Chisq) |
|---|---|---|---|---|---|---|---|---|
| **m0** | 3 | 39316 | 39338 | -19655 | 39310 | | | |
| **m1** | 4 | 39309 | 39339 | -19651 | 39301 | 8.42 | 1 | 0.004** |
| **m2** | 5 | 39311 | 39347 | -19650 | 39301 | 0.77 | 1 | 0.38 |
| **m3** | 6 | 39311 | 39355 | -19650 | 39299 | 1.40 | 1 | 0.24 |
| **m4** | **7** | **39305** | **39356** | **-19646** | **39291** | **8.27** | **1** | **0.004**** |
| **m4_1** | **8** | **39304** | **39362** | **-19644** | **39288** | **3.32** | **1** | **0.07** |
| **m5** | 8 | 39307 | 39365 | -19646 | 39291 | 0 | 0 | |
| **m6** | 9 | 39304 | 39369 | -19643 | 39286 | 5.45 | 1 | 0.02 * |
| **m4_2** | 9 | 39306 | 39371 | -19644 | 39288 | 0 | 0 | |
| **m7** | 10 | 39305 | 39378 | -19643 | 39285 | 2.18 | 1 | 0.14 |
| **m4_3** | 10 | 39308 | 39381 | -19644 | 39288 | 0 | 0 | |
| **m4_4** | 13 | 39308 | 39403 | -19641 | 39282 | 5.76 | 3 | 0.12 |
| **m4_5** | 14 | 39309 | 39411 | -19641 | 39281 | 0.70 | 1 | 0.40 |
| **m4_6** | 15 | 39310 | 39420 | -19640 | 39280 | 0.75 | 1 | 0.39 |
| Signif. codes: 0 '***' 0.001 '**' 0.01 '*' 0.05 '.' 0.1 ' ' 1 | | | | | | | | |

**Supplementary Table S4**. Comparison of multiple statistical models for the **(III) – delayed response**. The overview shows *npar*, which represents the number of parameters in each model, *AIC* (Akaike Information Criterion), *BIC* (Bayesian Information Criterion), *logLik* (Log-Likelihood), *deviance* which was used to assess the goodness of fit of a model, *Chisq* (Chi-Square),which represents the difference in deviance between the current model and the previous one, *Df* (Degrees of Freedom), which represents the degrees of freedom associated with the Chi-Square test and *Pr(>Chisq)* which represents p-values associated with the Chi-Square test. Model m4 (red) was the best fitting model and m4_7 (red) was the second-best fitting model.

### Results

**Linear mixed models (model m4: *StimIntFreq-LMM*) for each time window (I-III):**

| model | | | Anova | β and t-value | mean ± std |
|---|---|---|---|---|---|
| **(I) ON** | **Trial_m4** | | | | |
| | | real vs. sham | $\chi^2$ = 18.97, p < 0.001 | real-sham: β = 0.08 (SE = 0.02; t-value = 4.37, p < 0.001) | real (0.18±0.03) vs sham (0.1±0.03) |
| | | high vs. low Hz | $\chi^2$ = 10.88, p = 0.001 | high-low: β = 0.06 (SE = 0.02; t-value = 3.30, p = 0.001) | high (0.17±0.03) vs low (0.11±0.03) Hz |
| | | high vs. low mA | $\chi^2$ = 103.40, p < 0.001 | high-low: β = 0.2 (SE = 0.02; t-value = 10.17, p < 0.001) | high (0.23±0.03) vs low (0.04±0.03) mA |
| | **Mean_m4** | | | | |
| | | real vs. sham | $\chi^2$ = 11.29, p = 0.0007 | real-sham: β = 0.09 (SE = 0.03; t-value = 3.36, p = 0.0009) | real (0.18±0.03) vs sham (0.1±0.03) |
| | | high vs. low Hz | $\chi^2$ = 5.94, p = 0.01 | high-low: β = 0.06 (SE = 0.03; t-value = 2.44, p = 0.02) | high (0.17±0.03) vs low (0.10±0.03) Hz |
| | | high vs. low mA | $\chi^2$ = 46.92, p < 0.001 | high-low: β = 0.18 (SE = 0.03; t-value = 6.85, p < 0.001) | high (0.22±0.03) vs low (0.05±0.03) mA |
| **(II) OFF** | **Trial_m4** | | | | |
| | | real vs. sham | $\chi^2$ = 15.99, p < 0.001 | real-sham: β = 0.1 (SE = 0.03; t-value = 3.99, p < 0.001) | real (0.15±0.04) vs sham (0.04±0.04) |
| | | high vs. low Hz | $\chi^2$ = 2.92, p = 0.09 | high-low: β = 0.05 (SE = 0.03; t-value = 1.71, p = 0.09) | high (0.11±0.04) vs low (0.07±0.04) Hz |
| | | high vs. low mA | $\chi^2$ = 76.79, p < 0.001 | high-low: β = 0.2 (SE = 0.03; t-value = 8.76, p < 0.001) | high (0.21±0.04) vs low (-0.03±0.04) mA |
| | **Mean_m4** | | | | |
| | | real vs. sham | $\chi^2$ = 5.21, p = 0.02 | real-sham: β = 0.1 (SE = 0.04; t-value = 2.30, p = 0.02) | real (0.15±0.04) vs sham (0.05±0.04) |
| | | high vs. low Hz | $\chi^2$ = 0.75, p = 0.39 | high-low: β = 0.04 (SE = 0.04; t-value = 0.86, p = 0.39) | high (0.12±0.04) vs low (0.08±0.04) Hz |
| | | high vs. low mA | $\chi^2$ = 22.12, p < 0.001 | high-low: β = 0.20 (SE = 0.04; t-value = 4.70, p < 0.001) | high (0.20±0.04) vs low (-0.004±0.04) mA |
| **(III) OFF** | **Trial_m4** | | | | |
| | | real vs. sham | $\chi^2$ = 0.61, p = 0.43 | real-sham: β = 0.02 (SE = 0.03; t-value = 0.78, p = 0.43) | real (-0.04±0.02) vs sham (-0.06±0.02) |
| | | high vs. low Hz | $\chi^2$ = 1.52, p = 0.22 | high-low: β = 0.04 (SE = 0.03; t-value = 1.24, p = 0.22) | high (-0.03±0.02) vs low (-0.07±0.02) Hz |
| | | high vs. low mA | $\chi^2$ = 8.26, p = 0.004 | high-low: β = 0.08 (SE = 0.03; t-value = 2.88, p = 0.004) | high (-0.01±0.02) vs low (-0.09±0.02) mA |
| | **Mean_m4** | | | | |
| | | real vs. sham | $\chi^2$ = 0.57, p = 0.45 | real-sham: β = 0.02 (SE = 0.03; t-value = 0.76, p = 0.45) | real (-0.04±0.02) vs sham (-0.06±0.02) |
| | | high vs. low Hz | $\chi^2$ = 1.74, p = 0.19 | high-low: β = 0.04 (SE = 0.03; t-value = 1.32, p = 0.19) | high (-0.03±0.02) vs low (-0.07±0.02) Hz |
| | | high vs. low mA | $\chi^2$ = 9.19, p = 0.002 | high-low: β = 0.08 (SE = 0.03; t-value = 3.03, p = 0.002) | high (-0.01±0.02) vs low (-0.09±0.02) mA |

**Supplementary Table S5**. Distinct LMM (*pupil dilation ~ trials + stimulation + intensity + frequency + (1|ID))* for each time window (**(I)** the 3 sec during on stimulation, **(II)** the immediate response and **(III)** the delayed response) based on data at the level of individual trials (blue; Trial_m4) or on the average pupil dilation per session (grey; Mean_m4).

### Linear mixed models (model m4_1 *StimIntFreq-VAS-LMM*) for each time window (I-III)

| model | | | Anova | β and t-value | mean ± std |
|---|---|---|---|---|---|
| (I) ON | Trial_m4_1 | | | | |
| | | real vs. sham | $\chi^2$ = 2.69, p = 0.1 | real-sham: β = 0.03 (SE = 0.02; t-value = 1.64, p = 0.1) | real (0.16±0.03) vs sham (0.12±0.03) |
| | | high vs. low Hz | $\chi^2$ = 2.85, p = 0.1 | high-low: β = 0.03 (SE = 0.02; t-value = 1.69, p = 0.1) | high (0.16±0.02) vs low (0.12±0.03) Hz |
| | | high vs. low mA | $\chi^2$ = 28.79, p < 0.001 | high-low: β = 0.1 (SE = 0.02; t-value = 5.34, p < 0.001) | high (0.20±0.03) vs low (0.08±0.03) mA |
| | Mean_m4_1 | | | | |
| | | real vs. sham | $\chi^2$ = 2.35, p = 0.13 | real-sham: β = 0.04 (SE = 0.03; t-value = 1.53, p = 0.13) | real (0.15±0.03) vs sham (0.11±0.03) |
| | | high vs. low Hz | $\chi^2$ = 2.04, p = 0.15 | high-low: β = 0.04 (SE = 0.03; t-value = 1.43, p = 0.16) | high (0.15±0.03) vs low (0.11±0.03) Hz |
| | | high vs. low mA | $\chi^2$ = 14.69, p = 0.0001 | high-low: β = 0.11 (SE = 0.03; t-value = 3.83, p < 0.0002) | high (0.19±0.03) vs low (0.08±0.03) mA |
| (II) OFF | Trial_m4_1 | | | | |
| | | real vs. sham | $\chi^2$ = 2.42, p = 0.12 | real-sham: β = 0.04 (SE = 0.03; t-value = 1.56, p = 0.12) | real (0.12±0.04) vs sham (0.07±0.04) |
| | | high vs. low Hz | $\chi^2$ = 0.09, p = 0.76 | high-low: β = 0.008 (SE = 0.03; t-value = 0.30, p = 0.76) | high (0.1±0.04) vs low (0.09±0.04) Hz |
| | | high vs. low mA | $\chi^2$ = 20.10, p < 0.001 | high-low: β = 0.13 (SE = 0.03; t-value = 4.48, p < 0.001) | high (0.17±0.04) vs low (0.03±0.04) mA |
| | Mean_m4_1 | | | | |
| | | real vs. sham | $\chi^2$ = 0.95, p = 0.33 | real-sham: β = 0.04 (SE = 0.04; t-value = 0.98, p = 0.38) | real (0.12±0.04) vs sham (0.07±0.04) |
| | | high vs. low Hz | $\chi^2$ = 0.02, p = 0.90 | high-low: β = 0.006 (SE = 0.04; t-value = 0.13, p = 0.95) | high (0.1±0.04) vs low (0.1±0.04) Hz |
| | | high vs. low mA | $\chi^2$ = 6.43, p = 0.01 | high-low: β = 0.12 (SE = 0.05; t-value = 2.54, p = 0.01) | high (0.16±0.04) vs low (0.03±0.04) mA |
| (III) OFF | Trial_m4_1 | | | | |
| | | real vs. sham | $\chi^2$ = 0.009, p = 0.92 | real-sham: β = 0.003 (SE = 0.03; t-value = 0.09, p = 0.92) | real (-0.05±0.02) vs sham (-0.05±0.02) |
| | | high vs. low Hz | $\chi^2$ = 0.68, p = 0.41 | high-low: β = 0.02 (SE = 0.03; t-value = 0.82, p = 0.41) | high (-0.04±0.02) vs low (-0.06±0.02) Hz |
| | | high vs. low mA | $\chi^2$ = 2.76, p = 0.1 | high-low: β = 0.05 (SE = 0.03; t-value = 1.66, p = 0.1) | high (-0.02±0.02) vs low (-0.08±0.02) mA |
| | Mean_m4_1 | | | | |
| | | real vs. sham | $\chi^2$ = 0.003, p = 0.95 | real-sham: β = 0.002 (SE = 0.03; t-value = 0.06, p = 0.95) | real (-0.05±0.02) vs sham (-0.05±0.02) |
| | | high vs. low Hz | $\chi^2$ = 0.81, p = 0.37 | high-low: β = 0.02 (SE = 0.03; t-value = 0.90, p = 0.37) | high (-0.04±0.02) vs low (-0.06±0.02) Hz |
| | | high vs. low mA | $\chi^2$ = 3.18, p = 0.07 | high-low: β = 0.05 (SE = 0.03; t-value = 1.78, p = 0.08) | high (-0.02±0.02) vs low (-0.07±0.02) mA |

**Supplementary Table S6**. Distinct LMM for each time window (**(I)** the 3 sec during on stimulation, **(II)** the immediate response and **(III)** the delayed response) based on data at the level of individual trials (blue; Trial_m4_1 (*pupil dilation ~ trials + stimulation + intensity + frequency + VAS + (1|ID)*) or on the average pupil dilation per session (grey; Mean_m4_1 (*pupil dilation ~ stimulation + intensity + frequency + VAS + (1|ID)*)).

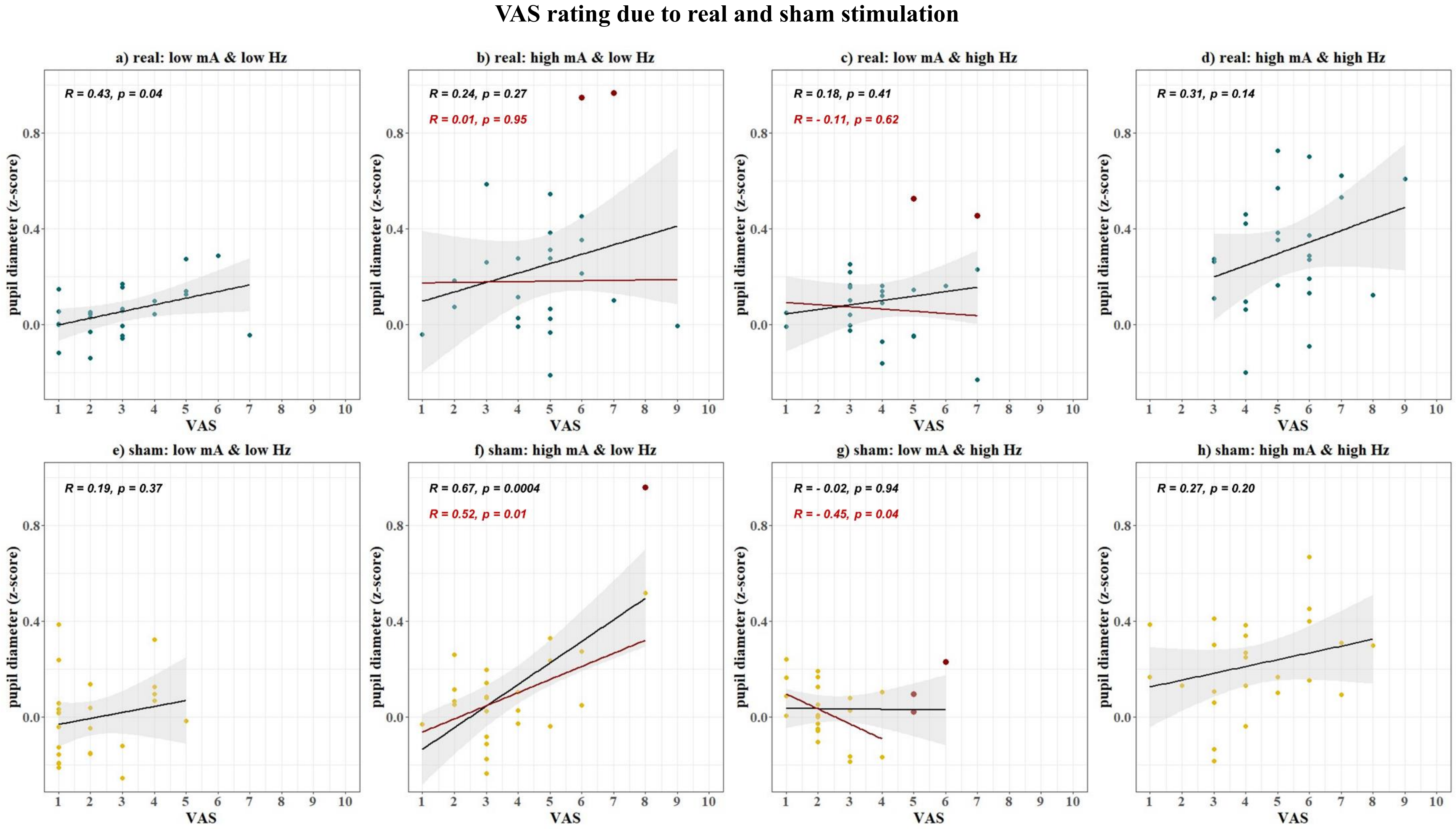


**Supplementary Figure S2**. Relationship between perception of sensations (VAS) due to stimulation and pupil dilation for (a-d) real stimulation (dark turquoise) and (e-h) sham stimulation (ochre). After each of the following sessions, a) real: low intensity & low frequency, b) real: high intensity & low frequency, c) real: low intensity & high frequency, d) real: high intensity & high frequency, e) sham: low intensity & low frequency, f) sham: high intensity & low frequency, g) sham: low intensity & high frequency, h) sham: high intensity & high frequency, VAS scores were recorded. For each correlation (a-h), a regression line (black) and a confidence interval (shadowed area) are given. Additionally, outliers (thicker red dots) were identified based on interquartile range and a corresponding regression line (red).

**Relationship between perception of sensations (VAS) due to stimulation and pupil dilation across subjects**

| | | R | p-value | CI | |
|---|---|---|---|---|---|
| | | | | lower | upper |
| **Real (R)** | | | | | |
| | **a) R: low mA low Hz** | **0.43** | **p = 0.04** | **0.03** | **0.71** |
| | b) R: high mA low Hz* | 0.01 | p = 0.95 | - 0.41 | 0.43 |
| | c) R: low mA high Hz* | - 0.11 | p = 0.62 | - 0.51 | 0.33 |
| | d) R: high mA high Hz | 0.31 | p = 0.14 | - 0.11 | 0.63 |
| **Sham (S)** | | | | | |
| | e) S: low mA low Hz | 0.19 | p = 0.37 | - 0.23 | 0.55 |
| | **f) S: high mA low Hz*** | **0.52** | **p = 0.01** | **0.14** | **0.77** |
| | **g) S: low mA high Hz*** | **- 0.45** | **p = 0.04** | **- 0.74** | **- 0.02** |
| | h) S: high mA high Hz | 0.27 | p = 0.20 | - 0.15 | 0.61 |

**Supplementary Table S7**. Overview of correlations between VAS rating after real (a-d) and sham stimulation (e-h). In addition to the significance (p-value), the confidence intervals (CI) are given for better evaluation and comparability of the different correlation effects. All correlations were adjusted for outliers (1.5*IQR); however, this adjustment occurred only for correlations marked with an asterisk. Significant correlations are bold.

**Relationship between subjective perception of sensations (VAS) of stimulation and pupil dilation across subjects.** To further investigate the potential effects of VAS on pupil dilation across subjects, VAS after each stimulation session and corresponding pupil dilation (averaged per subject across all trials within a stimulation condition) were correlated (see Supplementary Fig. S2, Supplementary Table S7). Correlations between VAS and pupil dilation considering outlier correction (see methods) were found for **a)** real stimulation: low intensity & low frequency (r = 0.43, p = 0.04), **f)** sham stimulation: high intensity and low frequency (r = 0.52, p = 0.01) as well as **g)** sham stimulation: low intensity & high frequency (r = - 0.45, p = 0.04). However, there were no correlation between VAS and pupil dilation for **b)** real stimulation: high intensity & low frequency (r = 0.24, p = 0.27), **c)** real stimulation: low intensity & high frequency (r = -0.11, p = 0.62), **d)** real stimulation: high intensity & high frequency (r = 0.31, p = 0.14), **e)** sham stimulation: low intensity & low frequency (r = 0.19, p = 0.37), **h)** sham stimulation: high intensity & high frequency (r = -0.45, p = 0.04). Thus, while higher mean pupil dilation was not consistently associated with higher VAS ratings across all subjects, three instances of significant associations between pupil dilation and VAS (two positive, one negative) were observed, suggesting that interindividual differences in subjective perceptions of stimulation also add variance to pupil ratings of stimulation effects.

**State of health**

The state of health queried in each case after stimulation to control for potential side effects are shown in Supp Table X. The following items were asked: (1) headache, (2) nausea, (3) tiredness, (4) dizziness, (5) tingling sensation at the previously stimulated area, (6) feeling of heat at the previously stimulated area, (7) reddening of the skin at the previously stimulated area, (8) skin irritation at the previously stimulated site, (9) impaired concentration, (10) itching at the previously stimulated area. Subjects indicated on a 4-point scale (0: not at all – 3: strong) to what extent they perceived potential side effects.

**Query of the state of health after the real and sham stimulation**

| | Real Stimulation | | | | Sham Stimulation | | | |
|---|---|---|---|---|---|---|---|---|
| | low Hz | | high Hz | | low Hz | | high Hz | |
| Item | low mA | high mA | low mA | high mA | low mA | high mA | low mA | high mA |
| 1 | 0.04 / 0.04 | 0.08 / 0.06 | 0 / 0 | 0 / 0 | 0 / 0 | 0 / 0 | 0 / 0 | 0 / 0 |
| 2 | 0.04 / 0.04 | 0 / 0 | 0 / 0 | 0 / 0 | 0 / 0 | 0 / 0 | 0 / 0 | 0 / 0 |
| 3 | 1.17 / 0.18 | 1.25 / 0.25 | 1.13 / 0.23 | 1.38 / 2.24 | 1 / 0.22 | 0.96 / 0.22 | 0.83 / 0.19 | 1.13 / 0.23 |
| 4 | 0 / 0 | 0 / 0 | 0 / 0 | 0 / 0 | 0 / 0 | 0.04 / 0.42 | 0 / 0 | 0 / 0 |
| 5 | 0.04 / 0.04 | 0.17 / 0.13 | 0.04 / 0.04 | 0.13 / 0.07 | 0 / 0 | 0.04 / 0.42 | 0 / 0 | 0.04 / 0.04 |
| 6 | 0 / 0 | 0.08 / 0.06 | 0.04 / 0.04 | 0.08 / 0.06 | 0 / 0 | 0.04 / 0.42 | 0 / 0 | 0.08 / 0.06 |
| 7 | 0 / 0 | 0 / 0 | 0 / 0 | 0 / 0 | 0 / 0 | 0 / 0 | 0 / 0 | 0 / 0 |
| 8 | 0 / 0 | 0 / 0 | 0 / 0 | 0 / 0 | 0 / 0 | 0 / 0 | 0 / 0 | 0 / 0 |
| 9 | 0.58 / 0.13 | 0.67 / 0.20 | 0.67 / 0.17 | 0.67 / 0.21 | 0.46 / 0.16 | 0.58 / 0.18 | 0.46 / 0.16 | 0.58 / 0.18 |
| 10 | 0 / 0 | 0 / 0 | 0 / 0 | 0 / 0 | 0.04 / 0.04 | 0.04 / 0.04 | 0 / 0 | 0 / 0 |

**Supplementary Table S8.** Query of the state of health (item 1-10) after real and sham stimulation. The mean value and standard deviation of the respective items (scale 0-3) across all subjects (N=24) for low vs. high frequency at given low vs. high intensity for real and sham stimulation are shown. No side effects occurred and the slightly higher score on item 3 (tiredness) is not due to the stimulation itself, but to the total duration of the experiment ( > 8h one day measurement).

**Perception of sensations (VAS)**
**at low vs. high frequency and intensity for real and sham stimulation**

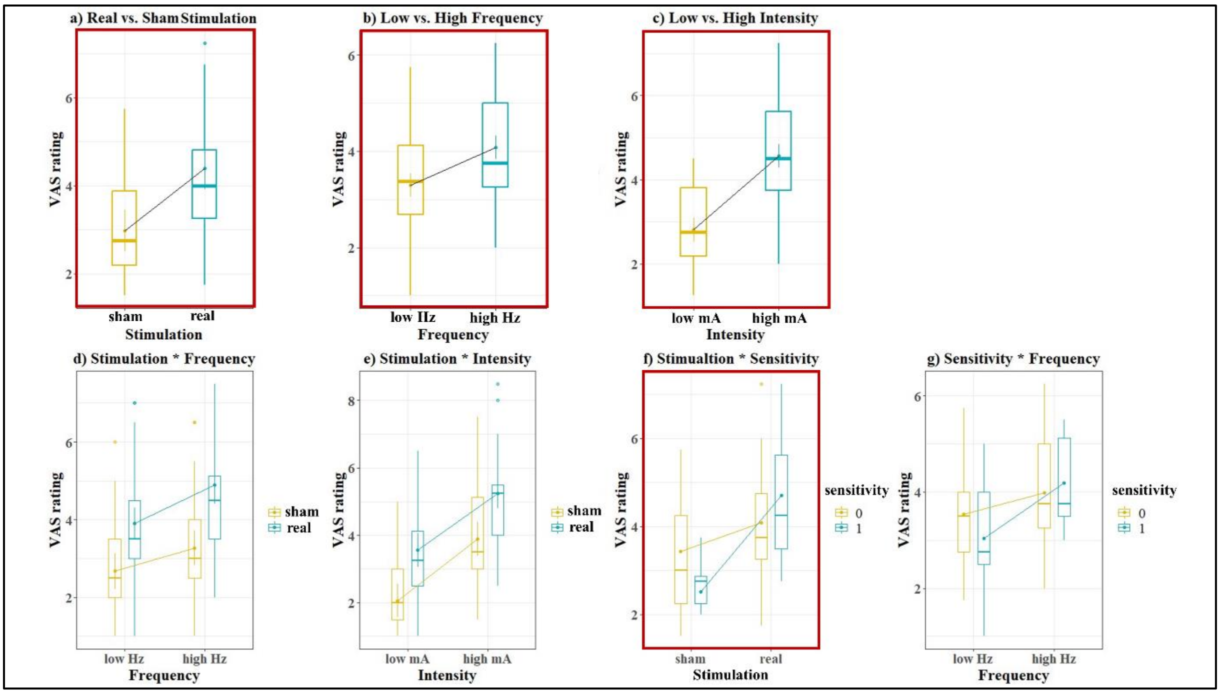


**Supplementary Figure S3**. Perception of sensations (VAS) (N =24) for a) real (turquoise) and sham (ochre) stimulation (F(1,20) = 17.85, p < 0.001), b) low (ochre) and high (turquoise) frequency (F(1,20) = 19.57, p < 0.001) and c) low (ochre) and high (turquoise) intensity (F(1,20) = 70.90, p < 0.001) as well as for d) Stimulation (real: turquoise; sham: ochre) * Frequency (F(1,20) = 4.06, p = 0.06), e) Stimulation (real: turquoise; sham: ochre) * Intensity (F(1,20) = 0.20, p = 0.66), f) Stimulation * Sensitivity (not sensitive (0): ochre; sensitive (1): turquoise) (F(1,20) = 4.86, p = 0.04) and g) Sensitivity (not sensitive (0): ochre; sensitive (1): turquoise) * Frequency (F(1,20) = 3.61, p = 0.07). The boxes outlined in red indicate significant main or interaction effects.

**Subjective higher perception of sensation (VAS) in response to stimulation.** The subjective perception of sensations (VAS) was not only higher for **(1)** *real stimulation* (M = 4.40, SD = 0.34) compared to sham stimulation (M = 2.97, SD = 0.29), (F(1,20) = 17.85, p < 0.001), but also for **(2)** *high frequency* (M = 4.08, SD = 0.28) compared to low frequency (M = 3.29, SD = 0.28), (F(1,20) = 19.57, p < 0.001), and **(3)** *high intensity* (M = 4.56, SD = 0.23) compared to low intensity (M = 2.81, SD = 0.23), (F(1,20) = 70.90, p < 0.001). There was no significant effect for either gender (F(1,20) = 0.02, p = 0.88), sporty (F(1,20) = 0.17, p = 0.69), or sensitivity (F(1,20) = 0.07, p = 0.79). There was a significant interaction between *sensitivity and stimulation* (F(1,20) = 4.86, p = 0.04), whereas sensitive subjects perceived higher sensations during real stimulation (M = 4.71, SD = 0.59) than during sham stimulation (M = 2.51, SD = 0.5); t (20) = 3.78, p = 0.006. Additionally there were trends for interactions between (1) stimulation and frequency (F(1,20) = 4.06, p = 0.06) as well as between (2) sensitivity and frequency (F(1,20) = 3.61, p = 0.07). However, there was no significant interaction between in stimulation and intensity (F(1,20) = 0.20, p = 0.66) (see Supplementary Fig. S3).

### Exploratory analysis of the interaction between VAS and stimulation and stimulation parameters.

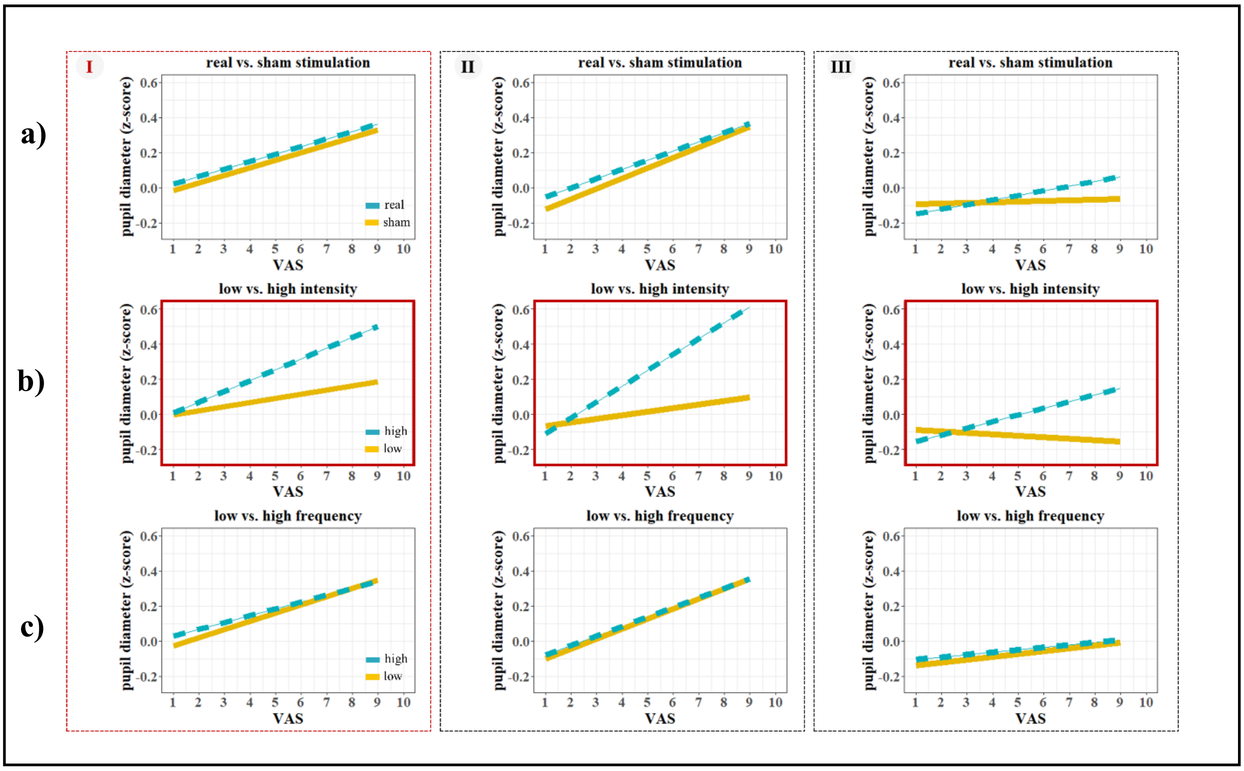


**Supplementary Figure S4**. Interaction plot between VAS and a) stimulation, b) intensity, and c) frequency for phasic (I) 'on stimulation', (II) 'immediate response', and (III) 'delayed response' (from left to right). Shown are estimated marginal means for different levels of VAS. The boxes outlined in red indicate the significant difference from between high and low frequency at all time windows (I-III): higher intensities and higher VAS rating predict a more dilated pupil than lower intensities ((I): $\chi^2$ = 12.08, p = 0.0005; (II) $\chi^2$ = 18.63, p < 0.001; (III) $\chi^2$ = 7.49, p = 0.006).

**Exploratory analysis of the interaction between VAS and stimulation and stimulation parameters.** Additional exploratory analysis based on the model

**<u>StimIntFreq*VAS-LMM</u>**

***pupil dilation* ~ trials + stimulation*VAS + intensity*VAS + frequency*VAS + sensitivity + (1|ID)***

**pupil dilation during (I) or (II) or (III)*

(Supplementary Fig. S4) showed for (**I**) 'on stimulation', that by adding VAS as individual interactions, while VAS was significant ($\chi^2$ = 38.74, p < 0.001), stimulation ($\chi^2$ = 3.39, p = 0.07) and frequency ($\chi^2$ = 3.38, p = 0.07) got marginal significant, while intensity ($\chi^2$ = 28.40, p < 0.001) still explained large portions of variance. There was no significant effect for sensitivity ($\chi^2$ = 0.12, p = 0.73). The model also revealed a significant interaction between VAS and intensity ($\chi^2$ = 12.08, p = 0.0005). Specifically, during low intensity an increase in VAS was associated with a 0.024 increase in pupil dilation, while during high intensity the effect was significantly larger with 0.062 increase in pupil dilation compared to low intensity stimulation (low vs. high: estimate -0.04, SE: 0.01; t (10466) = - 3.47, p = 0.0005). There was no significant interaction between VAS and stimulation ($\chi^2$ = 0.002, p = 0.1) nor VAS and frequency ($\chi^2$ = 0.58, p = 0.44).

The (**II**) 'immediate response' time-window showed, that by adding VAS as individual interactions, while VAS was significant ($\chi^2$ = 30.81, p < 0.001), intensity ($\chi^2$ = 19.93, p < 0.001) still explained large portions of variance, while stimulation ($\chi^2$ = 3.06, p = 0.08) and frequency ($\chi^2$ = 0.27, p = 0.60) were not or marginal significant. There was no significant effect for sensitivity ($\chi^2$ = 0.02, p = 0.88). The model also revealed a significant interaction between VAS and intensity ($\chi^2$ = 18.63, p < 0.001). Specifically, during low intensity an increase in VAS was associated with a 0.02 increase in pupil dilation, while during high intensity the effect was significantly larger with 0.1 increase in pupil dilation compared to low intensity stimulation (low vs. high: estimate -0.07, SE: 0.02; t (10535) = - 4.30, p < 0.001). There was no significant interaction between VAS and stimulation ($\chi^2$ = 0.15, p = 0.70) nor VAS and frequency ($\chi^2$ = 0.04, p = 0.85).

The (**III**) 'delayed response' time-window showed, that by adding VAS as individual interactions, neither VAS ($\chi^2$ = 3.40, p = 0.07), intensity ($\chi^2$ = 2.88, p = 0.09), stimulation ($\chi^2$ = 0.08, p = 0.78), frequency ($\chi^2$ = 0.90, p = 0.34) and sensitivity ($\chi^2$ = 0.29, p = 0.60) was significant. However, the model revealed a significant interaction between VAS and intensity ($\chi^2$ = 7.49, p = 0.006). Specifically, during low intensity an increase in VAS was associated with a -0.009 decrease in pupil dilation, while during high intensity the effect was significantly larger with 0.04 increase in pupil dilation compared to low intensity stimulation (low vs. high: estimate -0.05, SE: 0.02; t (7096) = - 2.73, p = 0.006). There was no significant interaction between VAS and stimulation ($\chi^2$ = 1.95, p = 0.16) nor VAS and frequency ($\chi^2$ = 0.03, p = 0.87).

**Supplementary Results 2**

**Additional analysis including random slopes for stimulation intensity:**

An additional analysis with random slopes in the model was added: ***pupil dilation* ~ trials + stimulation + intensity + frequency + VAS + (1+ intensity |ID)***

**pupil dilation during (**I**) or (**II**) or (**III**)*

During (**I**) 'on stimulation' the model revealed that VAS explained significant proportion of pupil variance ($\chi^2 = 33.27$, $p < 0.001$). Therefore, there was no increase in pupil dilation during real (M±SD: 0.16±0.02) as compared to sham (M±SD: 0.12±0.03) stimulation ($\chi^2 = 3.09$, $p = 0.08$) and during low (M±SD: 0.12±0.03) as compared to high (M±SD: 0.16±0.02) frequency ($\chi^2 = 3.13$, $p = 0.08$) explainable. However, pupil dilation was still increased during high (M±SD: 0.20±0.03) as compared to low (M±SD: 0.08±0.02) intensity ($\chi^2 = 15.71$, $p < 0.001$).

During (**II**) 'immediate response' the model revealed that VAS explained significant proportion of pupil variance ($\chi^2 = 27.96$, $p < 0.001$). Therefore, there was no increase in pupil dilation during real (M±SD: 0.12±0.04) as compared to sham (M±SD: 0.07±0.04) stimulation ($\chi^2 = 2.51$, $p = 0.11$) and during low (M±SD: 0.09±0.04) as compared to high (M±SD: 0.1±0.04) frequency ($\chi^2 = 0.1$, $p = 0.75$) explainable. However, pupil dilation was still increased during high (M±SD: 0.16±0.05) as compared to low (M±SD: 0.02±0.03) intensity ($\chi^2 = 6.33$, $p < 0.01$).

During (**III**) 'delayed response' the model revealed that VAS did not explain significant proportion of pupil variance ($\chi^2 = 2.67$, $p = 0.10$) anymore. There was also no increase in pupil dilation during real (M±SD: -0.05±0.02) as compared to sham (M±SD: -0.05±0.02) stimulation ($\chi^2 = 0.02$, $p = 0.89$) and during low (M±SD: -0.06±0.02) as compared to high (M±SD: -0.04±0.02) frequency ($\chi^2 = 0.76$, $p = 0.38$) explainable. Additionally, dilation was also not increased during high (M±SD: -0.02±0.02) as compared to low (M±SD: -0.08±0.02) intensity ($\chi^2 = 2.55$, $p = 0.11$).

### Changes in pupil dilation during real and sham stimulation for low and high stimulation intensity and frequency based on *StimIntFreq-LMM* and *StimIntFreq-VAS-LMM*

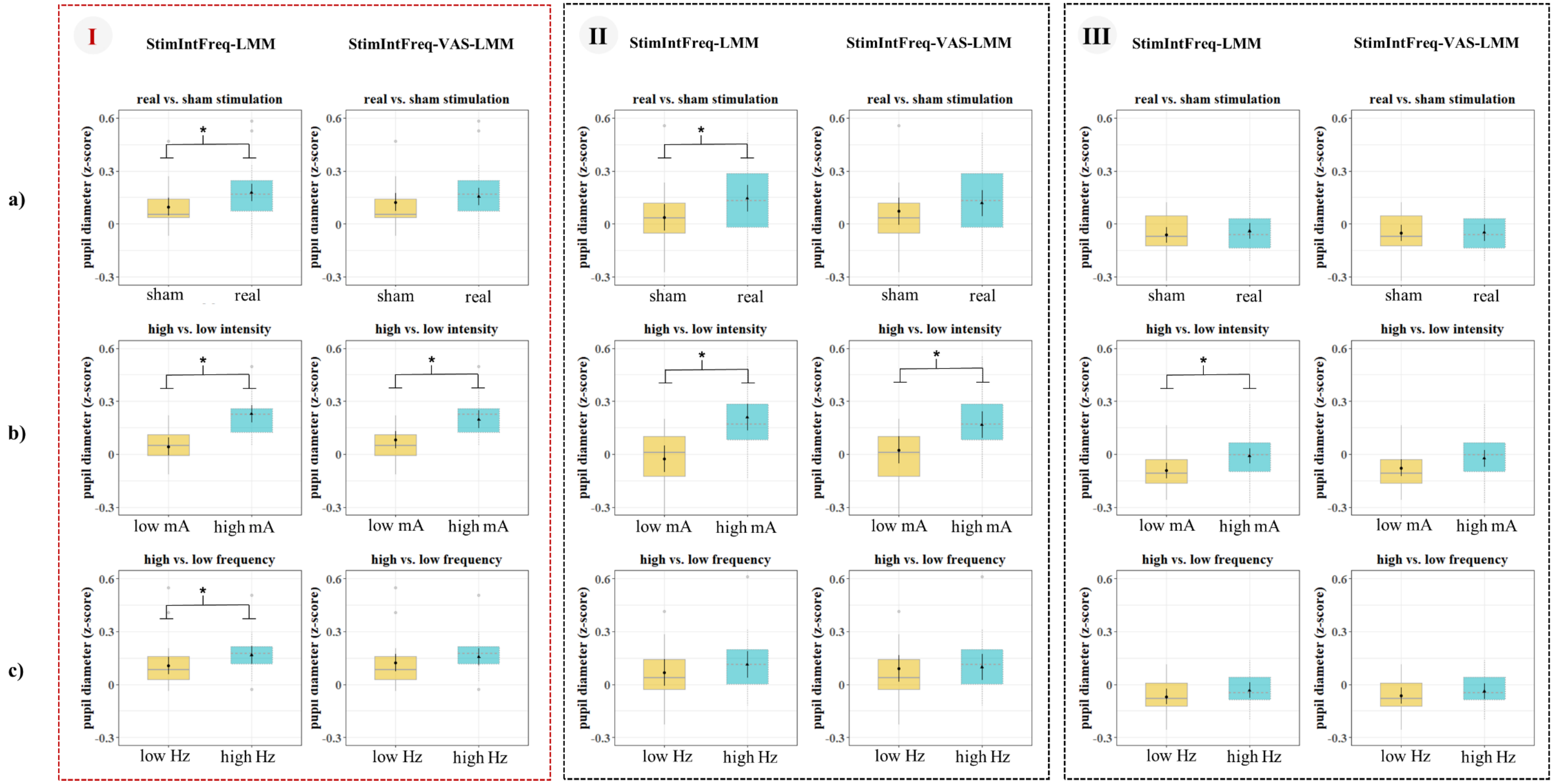


**Supplementary Figure S5**. Pupil diameters for **a)** real (turquoise) and sham (ochre) stimulation, **b)** high (turquoise) and low (ochre) intensity and **c)** high (turquoise) and low (ochre) frequency. The dashed vertical red lines indicate the time window when the **stimulation** was (I) **on**, the (II) **immediate response** following to the first dashed vertical black line and the subsequent (III) **delayed response** to the second dashed vertical black line. The boxplots for the individual time points are based on ***StimIntFreq-LMM and StimIntFreq-VAS-LMM***, the asterisks indicate significant differences between conditions.
